# Myosin-5 varies its steps along the irregular F-actin track

**DOI:** 10.1101/2023.07.16.549178

**Authors:** Adam Fineberg, Yasuharu Takagi, Kavitha Thirumurugan, Joanna Andrecka, Neil Billington, Gavin Young, Daniel Cole, Stan A. Burgess, Alistair P. Curd, John A. Hammer, James R. Sellers, Philipp Kukura, Peter J. Knight

**Author notes:** Correspondence: Yasuharu Takagi, Philipp Kukura, Peter J. Knight. Structural Biology Lab, Pearl Research Park, SBST, Vellore Institute of Technology, Vellore-632 014, India. Human Technopole, Viale Rita Levi-Montalcini 1, 20157, Milan, Italy (Joanna Andrecka). Department of Biochemistry and Molecular Medicine, West Virginia University, Morgantown, WV, U.S.A. Refeyn Ltd., Unit 9, Trade City, Sandy Ln W, Littlemore, Oxford OX4 6FF, U.K.

## Abstract

Molecular motors employ chemical energy to generate unidirectional mechanical output against a track. By contrast to the majority of macroscopic machines, they need to navigate a chaotic cellular environment, potential disorder in the track and Brownian motion. Nevertheless, decades of nanometer-precise optical studies suggest that myosin-5a, one of the prototypical molecular motors, takes uniform steps spanning 13 subunits (36 nm) along its F-actin track. Here, we use high-resolution interferometric scattering (iSCAT) microscopy to reveal that myosin takes strides spanning 22 to 34 actin subunits, despite walking straight along the helical actin filament. We show that cumulative angular disorder in F-actin accounts for the observed proportion of each stride length, akin to crossing a river on variably-spaced stepping stones. Electron microscopy revealed the structure of the stepping molecule. Our results indicate that both motor and track are soft materials that can adapt to function in complex cellular conditions.

## Introduction

Myosin-5a is a vital cellular motor ^1–3^. It transports cargoes by converting chemical energy from ATP hydrolysis into unidirectional, processive motion along an actin filament. It has two heads, each comprising a motor domain connected to a coiled-coil tail via a long, *α*-helical lever that extends from the converter subdomain of the motor ^4–6^ (Figure 1A). The molecule terminates in a dimeric domain that binds cargo. The ~22 nm lever helix is stabilized by six calmodulin-family light chains ^6, 7^. The mechanochemical mechanism is tightly coupled to ATP hydrolysis and the kinetics are limited by ADP release ^8–10^. ATP binding induces a change in motor conformation that allows catalysis of ATP hydrolysis and is coupled to a swing of the lever into a ‘primed’ (also known as pre-powerstroke) orientation. Stereospecific binding of this primed motor to F-actin catalyzes release of phosphate and ADP, coupled to swinging of the primed lever (the powerstroke) back to its unprimed (post-powerstroke) angle, thus causing movement of attached cargoes along the actin ^5–7, 11–14^. Binding of another ATP causes rapid dissociation from actin and the cycle repeats. The ATPase kinetics of the two-headed myosin-5a on actin are such that both motor domains are bound to actin most of the time, with ADP in the active site ^10^. In the two head-attached state, ADP release from the lead head is markedly suppressed (‘gated’) allowing the trail head to release its ADP and then rapidly bind ATP resulting in dissociation of this head from actin ^9,10, 15, 16^. The attached head then undergoes the powerstroke that repositions the unattached head to search for a new forward binding site on actin ^16, 17^. This process repeats, resulting in multiple ATP-dependent steps on actin per diffusional encounter of a myosin-5a molecule ^10, 15, 18^. Despite this consensus, it is controversial what size steps myosin-5a takes along actin, what role actin structure plays in the pattern of myosin-5a movement and what is the structural basis of gating of ADP release.

**Figure 1.**
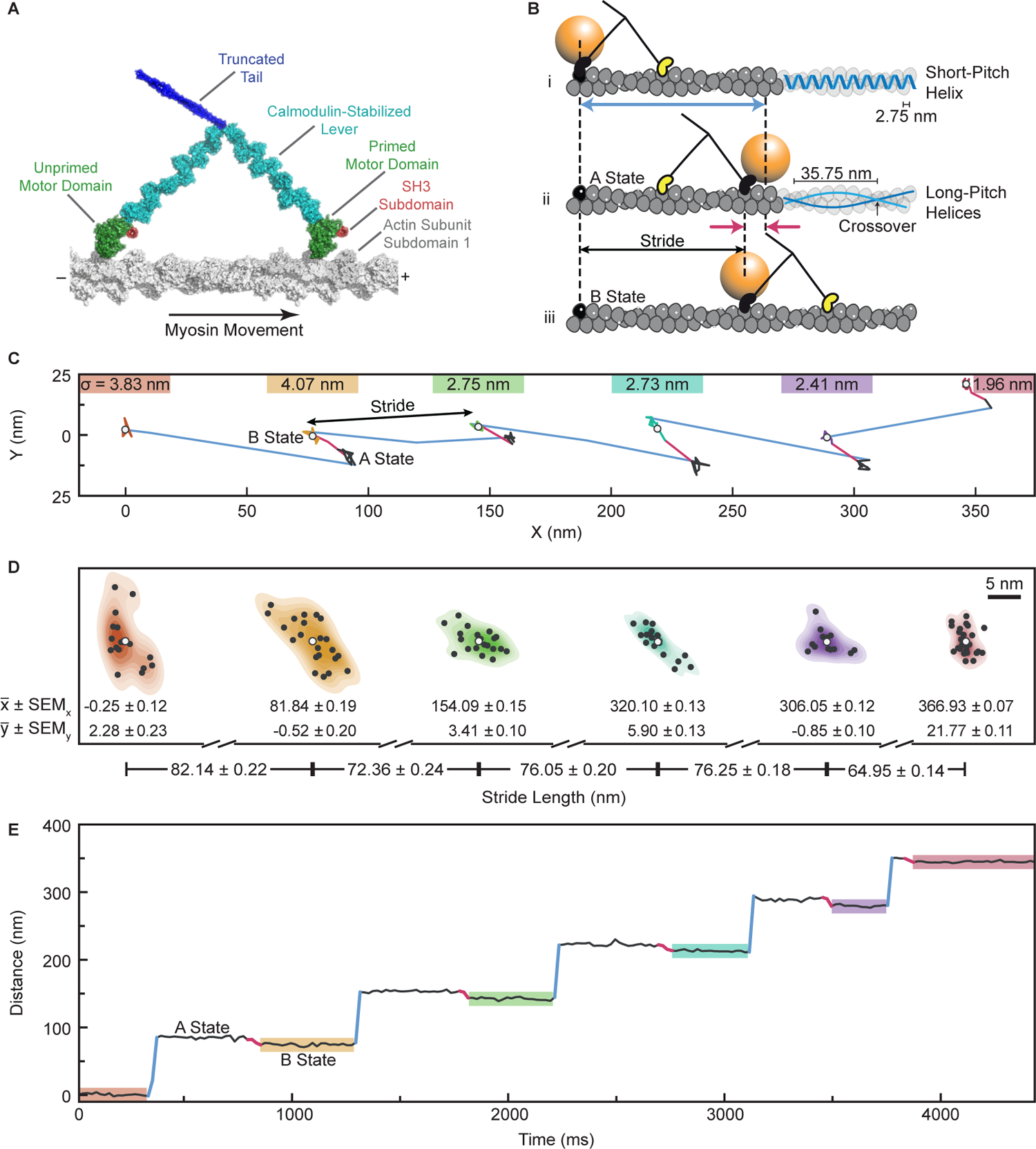
Trajectory along F-actin of a myosin-5a motor labelled with a gold bead. (A) Diagram of myosin-5a, truncated after the proximal coiled coil. (B) Schematic of two consecutive myosin steps, depicting first, the labelled head movement (blue arrow; B-state to A-state) and second, the AB transition of the bead (red arrows) accompanying the unlabeled head movement and allowing measurement of the labelled head stride (B-state to B-state). Also shown at right is actin’s short-pitch helix (top filament; bar indicates separation of successive subunits along that helix) and two long-pitch helices (middle filament; bar indicates spacing of successive crossovers). (C) The XY trace of a typical run along F-actin of a gold labelled myosin-5a molecule, highlighting labelled head strides (blue) and AB transitions (red). We report stride lengths as the Euclidean distance between the mean x and y values of consecutive B states. Localization precisions (σ), defined as the root sum of squares of x and y standard deviations of stationary states, marked above each B state. (D) The 2D kernel density estimates of bead positions highlighted in (C) with black points at each localization and white points at the mean values (x, y). Distances between the means demonstrate our high precision of stride length measurements. Uncertainty on mean particle position is standard error on mean (SEMx, SEMy), propagated to the error on stride sizes. (E) Distance along the filament versus Time trace of the XY plot in (C), again, highlighting labelled head movements (blue) and AB transitions (red).

F-actin structure dictates the options for stepping by myosin-5a. It is a polar, helical polymer of globular actin subunits in which a single, left-handed, short-pitch helix is formed by successive subunits having an axial separation of 2.75 nm and a more variable left-handed rotation of −166° ^19, 20^. In side-view, this gives an appearance of two coaxial, right-handed helices of subunits, staggered by a half subunit, that cross over one another every ~36 nm (Figure 1B). For myosin-5a to walk straight, i.e., in a single azimuthal plane, along F-actin it must attach to a sequence of subunits of similar azimuth. These will generally be 13 subunits (35.75 nm) apart along the filament axis, because 13 x −166° is close to an integer number of turns (6 turns) of the short-pitch helix. When the trailing motor domain of a straight-walking molecule lets go, moves past the leading motor, and reattaches, the ‘stride’ that that motor takes is thus typically 26 subunits (71.50 nm), with each motor making 13-subunit steps ahead of its partner motor and the whole molecule making a forward movement of 13 subunits (35.75 nm). Myosin-5a can take strides of this length because of its long levers. The inherent polarity of the actin filament ^21^ dictates the direction of movement of the myosin.

The structure of myosin-5a bound to actin by both motors while walking slowly in the presence of rate-limiting, micromolar ATP was captured by negative-stain electron microscopy (nsEM) ^6^. The trailing head lever angle resembled the well-known ‘rigor’ complex formed by myosin heads on actin in the absence of ATP. By contrast the lead head lever emerged from the rear side of the leading motor domain and pointed backwards to meet the trailing head lever at their junction with the tail. The lead motor appeared to be in a near-primed position expected of a motor at the start of its working stroke, whereas the trail head converter was at the end of the working stroke ^7^. However, this interpretation has not been universally adopted, in part because the measured rapid release of phosphate from the lead motor is associated with a motor transition to the unprimed state, and partly because of concerns that nsEM may have produced an artifactual structure. We have therefore used cryogenic electron microscopy (cryoEM) to reinvestigate the structure of the walking molecule. CryoEM avoids many possibilities for artifacts that are possible for nsEM because the sample is flash-frozen during activity without adsorption to a carbon film, fixation, staining or drying ^22, 23^, but it has not previously been successful for studying myosin-5a walking on actin.

Evidence suggests that myosin-5a must vary its stride length in order to walk straight. Recent high-resolution studies of F-actin ^19, 20, 24, 25^ find a subunit rotation of −166.5°, which implies a −4.5° (left-handed) movement around the actin filament per 13-subunit step. A run of 20 steps of 13 subunits, covering 715 nm, would thus entail a 90° rotation around the actin. Furthermore, nsEM of F-actin shows variable spacing of crossovers along each filament, which has been interpreted as arising from random variation in the rotation angle between successive subunits with a standard deviation of 6° ^26–28^. This cumulative angular disorder (CAD) means that the 13^th^ subunit will not always be best oriented to allow straight walking, so a variable stride would be necessary. Although CAD was demonstrated in early cryoEM studies of F-actin ^29, 30^, recent cryoEM studies have not commented on the presence of CAD in their specimens, leaving open to question whether CAD was merely an artifact of earlier preparation protocols. Although CAD in F-actin has been characterized, it has been unclear whether it has significance in the cell or is simply a biophysical curiosity. We have therefore analyzed recent cryoEM data of F-actin ^19^ for evidence of CAD and find that it does indeed exist and has a major impact on myosin-5a stepping behavior.

It has been controversial whether myosin-5a takes uniform or variable steps as it walks along F-actin. nsEM indicated that the two heads of myosin-5a steps were commonly spaced 13 actin subunits apart but spacings of 11 and 15 actin subunits were also frequent ^6,31^. In contrast, optical trapping or single molecule fluorescence assays of active molecules have shown a broad, continuous distribution of movements with a mean of 37 nm or 75 nm depending on the site and number of labels per molecule ^10, 32–37^. Recently, we used interferometric scattering (iSCAT) microscopy ^38–40^ to study myosin-5a striding on actin ^17^. A 20-nm gold nanoparticle coupled to the N-terminus of one motor (Figure 1B) enables precise localization (~1 nm) and high temporal resolution, in the order of milliseconds. The initial analysis of stride lengths again showed a single broad peak at 74 nm, consistent with previous studies. All these assays of active molecules have allowed a conclusion that all steps are 13 actin subunits, with a Gaussian broadening resulting from experimental and measurement uncertainties ^13–15, 32–34^. However, it was puzzling why the iSCAT peak was so broad given the method’s inherently high spatial resolution. We have therefore explored alternative methods of iSCAT data analysis and discovered that the breadth can be resolved into a family of discrete stride lengths.

The iSCAT study also detected a shift in position of the gold bead when the unlabeled head made its forward stride. This ‘AB transition’ is linked to the working stroke of the labelled motor that carries the unlabeled motor towards its next binding site (Figure 1B). It allows quantification of the dwell times of both heads of the molecule, and this showed that the kinetics of the labeled and unlabeled heads were indistinguishable. In our new analysis we exploit this phenomenon to yield a fuller understanding of myosin-5a stepping.

Here we study myosin-5a during its walk along actin filaments by combining cryoEM, nsEM and iSCAT microscopy. We show that myosin-5a is an inherently imprecise stepper. It exhibits multimodal step- and stride-length distributions that were not resolved by previous, lower resolution, single molecule stepping studies. We further show that single molecules of myosin-5a can walk straight, varying each step length to accommodate the variability in actin filament structure that is caused by CAD and thus do not exhibit step by step variations in their azimuthal position on the actin filament. The new EM data are consistent with our earlier proposal that a preferred angle between the two levers places the motors 36 nm apart along actin ^17^, with an additional, thermally-driven elastic variation of this angle and in head shape allowing variable step size.

## Results

### iSCAT resolves varying stride lengths of myosin-5a

Molecules of myosin-5a construct labelled on the N-terminus of one motor with a 20-nm gold bead were tracked by iSCAT microscopy at 100 Hz as they walked along actin in the presence of 10 µM ATP. For each molecule, the x, y position of the bead was determined with sub-nanometer precision (0.9 nm localization precision) ^17^ in each frame, allowing the stride and the AB transition to be identified (Figure 1C and E). The difference between mean positions of the bead in successive B-states yields the length of each stride taken by the labelled head (Figure 1B - D).

The histogram of measured stride lengths is strikingly multimodal, with very little overlap between adjacent peaks (Figure 2A). This demonstrates unequivocally that myosin-5a takes strides of varying but quantized length and that iSCAT microscopy has the spatial precision to resolve these. The separation of adjacent peaks of 7 Gaussians fitted to the binned data is 5.45 ± 0.40 nm (mean ± SD), which corresponds to a separation of two actin subunits along the filament (5.5 nm), that is, it corresponds to neighboring subunits along one of the two long-pitch strands (Figure 1B). The major peak at 71.8 nm corresponds to a distance spanning 26 actin subunits (expected value, 71.5 nm). This is the stride length expected from two canonical 13-subunit steps. The other peaks therefore correspond to strides that land on subunits nearby to the 26^th^ along the same long-pitch actin strand. The family of peaks thus correspond to strides spanning 22, 24, 26, 28, 30, 32 and 34 actin subunits. Using the average of all the peaks, weighted by their normalized area, the spacing of subunits along F-actin is 2.758 ± 0.016 nm. Since the myosin motor binds stereo-specifically to actin ^41^ the width of each peak likely arises from imprecision of measurement. This is expected to be independent of stride length and indeed an excellent fit to the data is obtained using a single width parameter (standard deviation (SD) = 1.195 nm) for all 7 peaks (Figure 2A). Using iSCAT microscopy, it is therefore possible to describe any individual processive run by the precise numbers of actin subunits traversed in each of the strides. For example, the run shown in Figure 1C-E can be described as 30, 26, 28, 28, 24.

**Figure 2.**
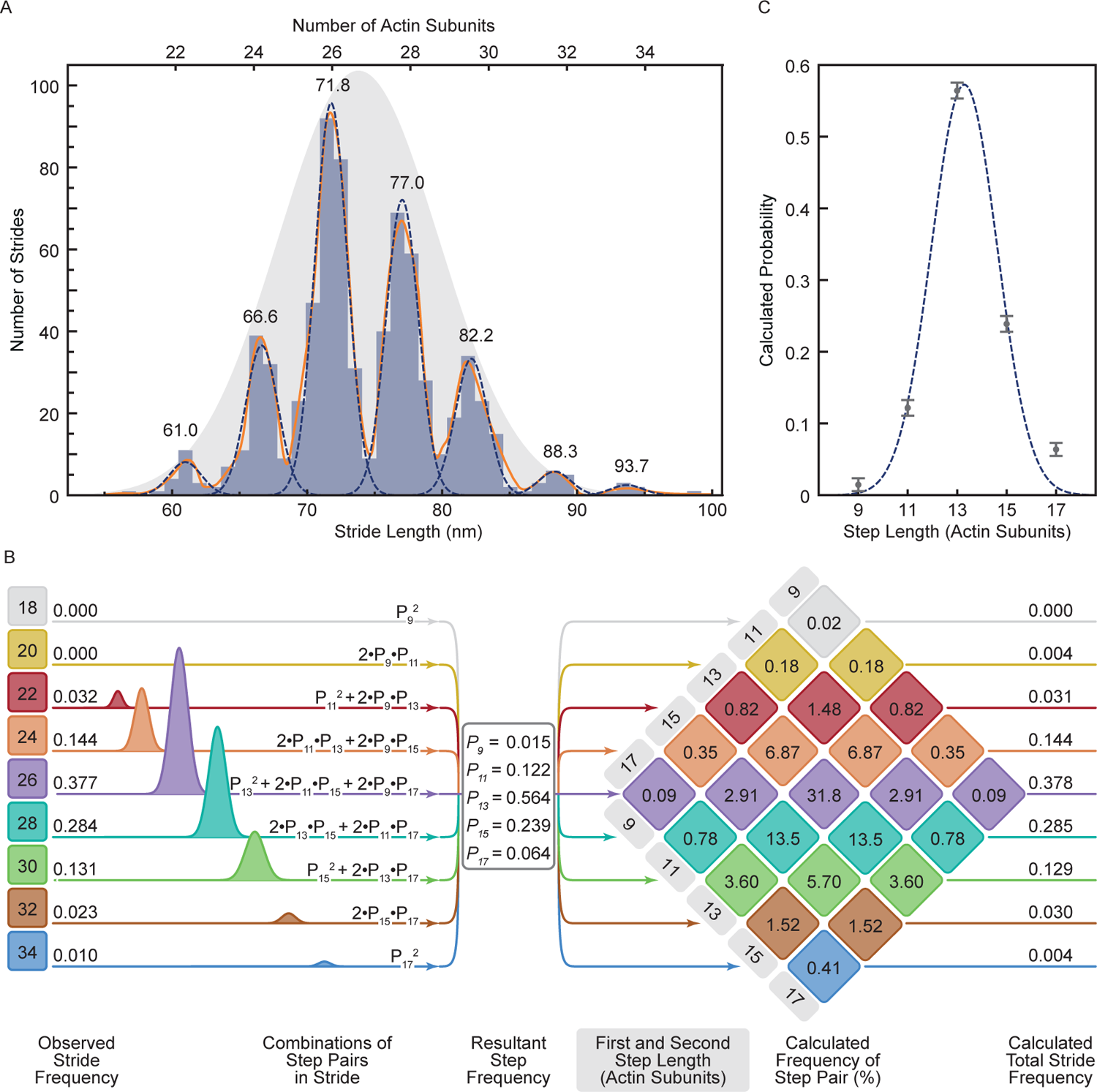
Analysis of strides taken by myosin-5a on F-actin. (A) Histogram of measured myosin-5 strides (N = 96 molecules, 725 strides). A Gaussian kernel density estimate, with locally adaptive bandwidth, is shown (solid orange line), as well as seven individual Gaussians (dashed lines). Mean stride length (nm) of each Gaussian is labelled above the fits, and calculated number of actin subunits traversed marked. The single Gaussian that expresses the overall mean and SD of the dataset is shown (grey shaded), rescaled to show fit to relative frequencies of the strides. (B) A schematic depicting the set of 9 equations used to calculate the probability of a given step length (Px, where x = 9, 11, 13, 15, 17 actin subunits) from the areas of the Gaussian fits to the stride peaks listed on the left side. Center panel lists the calculated step length probabilities. On the right is the multiplication diamond using the calculated probabilities of each step length to determine the percentage frequency of a stride resulting from each combination of step lengths. Combinations of step lengths that result in the same stride length are of the same color. The total predicted frequencies of each stride length are listed on the right for comparison with the observed frequencies listed on the left. (C) Step probabilities with fitted Gaussian.

The data indicate that all molecules behaved similarly. Thus, comparison of the trajectories of the 96 molecules (Figure S2A) shows that all molecules take a range of stride lengths, rather than some being inherent short-striders and others, long-striders. Also, the distribution of stride lengths is independent of the number of strides taken in a processive run (Figure S2B). Furthermore, a *χ*^2^ analysis showed that the length of each stride was independent of the length of the preceding stride. These are all features to be expected if successive strides are independent events. The dataset of strides has an overall mean of 73.7 nm and SD of 5.66 nm. These overall values are close to those obtained in earlier studies using other methods that did not resolve the component peaks ^32–34, 37^. This indicates that the flexibility in stepping that we observe by iSCAT was also present in all those studies. This mean and SD can be represented by a Gaussian distribution and scaling its amplitude shows that it is a good fit to the relative amplitudes of the component peaks (Figure 2A). This indicates that the underlying cause of variable stride length has a Gaussian character. We investigate this further below.

It is important to note that the overall mean value of stride length falls between our observed peaks, and thus does not correspond to a stride length that is ever taken. This is because binding sites for motors on F-actin are not separated by this mean value. By using high resolution iSCAT microscopy, we have thus revealed that molecules of myosin-5a take strides of variable length during a single processive run.

### Myosin-5a step frequencies can be estimated from the stride frequencies

Quantitative analysis of the relative frequencies of each stride length (Figure 2B) reveals further insights into the stepping behavior of myosin-5a. Each stride length is the sum of a forward step by first the unlabeled motor and then the labelled motor. For example, a 24-subunit stride can result from 4 different combinations (1^st^ step, 2^nd^ step): (11, 13), (13, 11), (9,15), and (15,9) actin subunits (Figure 2B). Using the measured relative frequencies of the 7 stride lengths (areas under the fitted peaks; Figure 2A - B), we found the least squares best fit solution to the set of simultaneous equations containing the probabilities of the 5 underlying step lengths (motors spanning 9, 11, 13, 15, and 17-subunits) (Figure 2B, left panel). The fit was robust and excellent, with only small differences between the observed frequencies and those back-calculated from the fitted values of step probabilities (Figure 2B, right panel). The values of step probabilities versus step length are well fitted by a single Gaussian of mean 13.3 with SD 1.32 actin subunits (with associated fitting errors, 0.12 and 0.10 actin subunits, respectively) (Figure 2C). These properties are strong evidence that the walking process is the result of a random and independent selection of step lengths governed by an underlying Gaussian probability distribution.

The range of step sizes that is implicit in the multiple stride lengths shows that the canonical 13-subunit step is not as abundant as previously thought. Thus, only 56% of all steps are 13-subunits (Figure 2B, middle panel, and Table 1). From the calculated relative abundance of the various step lengths (Figure 2B), the average step length is 13.469 actin subunits (= 37.04 nm) (Table 1). Likewise, only 38% of the strides are 26-subunits, and this includes combinations of 11+15 subunit steps in addition to two 13-subunit steps. Only 32% of all strides are a combination of two consecutive 13-subunit steps (Figure 2B, right panel).

**Table 1.** Calculated probabilities of each step length from iSCAT, cryoEM, and nsEM imaging. The mean step length and standard deviation in step lengths are also tabulated.

| Step Length<br>(actin subunits (asu)) | Step Probability |  |  |
| --- | --- | --- | --- |
|  | iSCAT | cryoEM | nsEM |
| 9 | 0.014 |  | 0.025 |
| 11 | 0.122 | 0.106 | 0.081 |
| 13 | 0.564 | 0.770 | 0.754 |
| 15 | 0.239 | 0.125 | 0.127 |
| 17 | 0.064 |  | 0.013 |
|  | iSCAT | cryoEM | nsEM |
| Mean Step (asu) | 13.469 | 13.115 | 13.044 |
| SD (asu) | 1.585 | 1.035 | 1.196 |
| Mean Step (nm) | 37.039 | 36.066 | 35.871 |
| SD (nm) | 4.357 | 2.846 | 3.289 |

This analysis to determine step frequencies shows that myosin-5a is a more flexible stepper than has been widely recognized, and that the widely stated 36 nm step accounts for only about half of the steps taken.

### Structure of myosin-5a on actin using cryoEM

We used cryoEM for the first time to examine the structure of full-length myosin-5a during stepping on actin, to test the earlier conclusions from nsEM. In practice, it proved difficult to get these samples into the holes of the support film, but a small dataset has been obtained that delineates the structures present. In the presence of ATP, low calcium concentration and no cargo adaptor molecules, myosin-5a is mainly compactly folded ^43–46^. These folded molecules are autoinhibited and bind only weakly to actin ^46^. However a small fraction of molecules is found to be active and to move processively along actin ^46^, so these should be present in samples flash frozen and examined by cryoEM. 55% of the myosin molecules observed were in the extended conformation and attached by both heads to the same actin filament (Figure S3). This abundance of myosin-5a molecules attached to actin is not surprising because we used a high ratio of myosin to actin (1 molecule per 2.5 actin subunits). Counts of myosin molecules per µm actin suggest that less than 10% were attached, which is consistent with the typical ~10-fold regulation of actin-activated ATPase activity in full-length myosin-5a preparations ^44^. The relative scarcity of detached molecules may arise from their adsorption to the carbon film.

In raw cryoEM images, the motor domains, levers and the first, coiled-coil segment of the tail are frequently identifiable (Figure 3A). The two heads of doubly-attached molecules typically show asymmetry recalling the appearance seen in negative stain 6, with the two levers pointing in opposite directions. A global average of doubly-attached molecules with motor domains 13 actin subunits apart improves clarity (Figure 3D). Both motor domains are attached on the leading side of sub-domain 1 of actin (compare with atomic model, Figure 3B), with the N-terminal SH3-fold sub-domain extending as far forward as the axial position of sub-domain 1 of the next actin subunit, as in models of primed and unprimed heads on actin (Figure 3B) ^47^. There is lower density between the motor and lever in the trail head than in the lead head, again in accord with the expectations from molecular models. The levers are visible throughout their length and unite at the head-tail junction without a gap, showing that the two polypeptide chains of the proximal coiled coil tail are not stretched apart by the stress within the doubly-attached molecule. All these features recall those seen using nsEM ^6, 31^.

**Figure 3.**
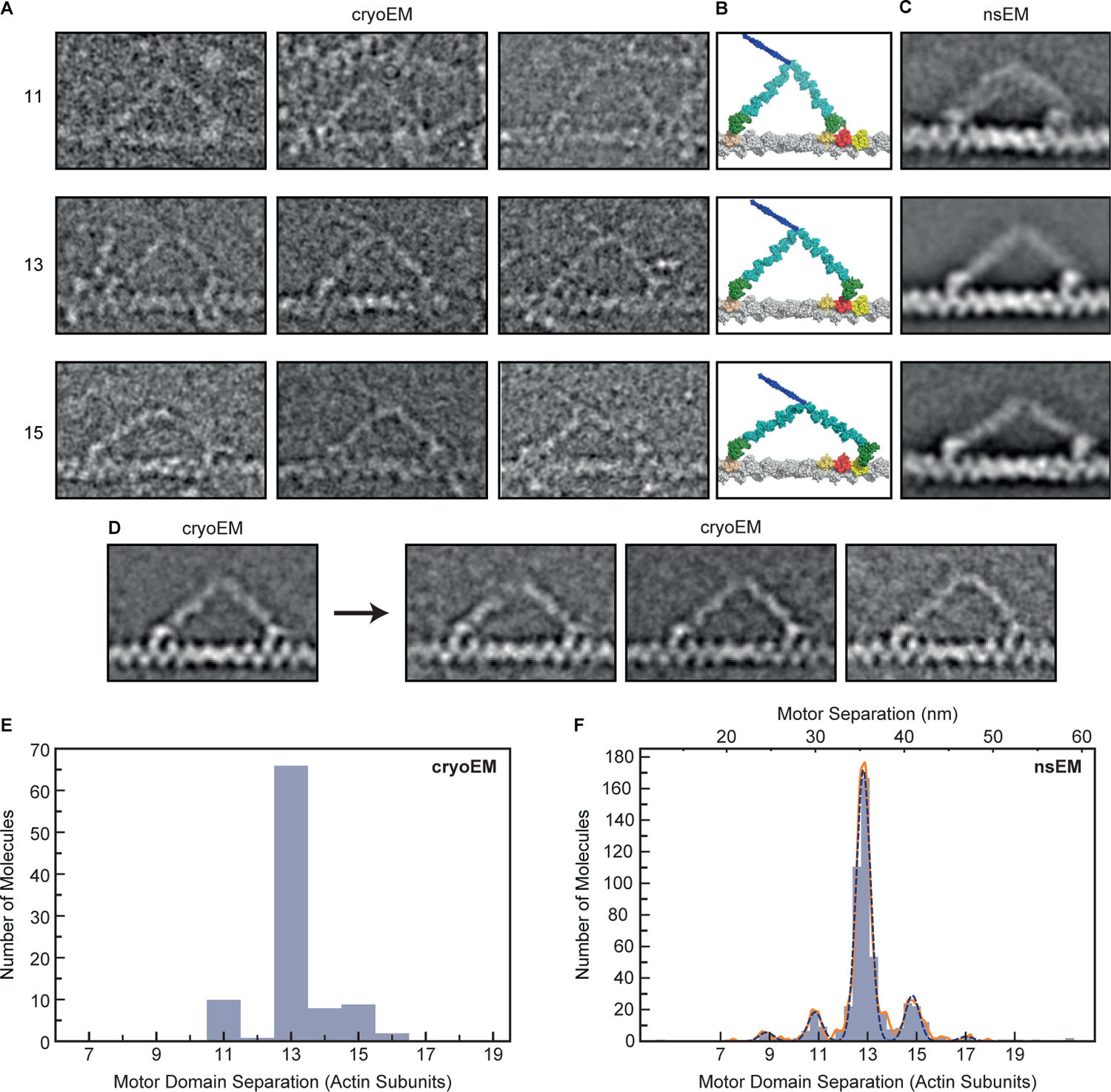
CryoEM and nsEM of myosin-5a walking on F-actin. (A) Gallery of single cryoEM images depicting myosin-5 molecules with step lengths of 11, 13 or 15 actin subunits. Contrast inverted (protein white). (B) Atomic models corresponding to (A), constructed according to Vale and Milligan ^42^, assuming a 13/6 helix for F-actin. Actin subunits 11, 13 or 15 subunits away from the trailing motor domain are colored pink, red and yellow respectively. (C) Averaged nsEM images of myosin-5 molecules with step lengths of 11, 13 or 15 actin subunits. Note that for the step length of 11 subunits, the leading motor is in an unprimed conformation, unlike for 13 and 15 subunit steps. (D) Left panel: averaged cryoEM image of myosin-5 molecules with step length of 13 actin subunits. Right panels: result of classification into three subclasses based on features in the leading lever. Note that the right subclass has a convex leading lever shape, but the lever still emerges from the trailing side of the motor domain. (A-D) Actin barbed (+) end and myosin-5a leading head are on the right in all panels. All panels (A-D) are scaled to match the panels in (B), in which 13 actin subunits span a distance of 35.75 nm. (E) Measured separation between motor domains from cryoEM data. (F) Measured separation between motor domains from nsEM data. A Gaussian kernel density estimate, with locally adaptive bandwidth, is shown (solid orange line), as well as five individual Gaussians (dashed blue lines). The number of actin subunits is shown along the bottom, and the corresponding separation distance in nanometers along the top of the figure.

In the trailing head the lever emerges from the leading side of the motor, near the SH3 domain, as expected for an unprimed head. The angle between the trailing lever and F-actin axis in the global average where the two motor domains are 13 subunits apart is 40°, as previously found by nsEM at rate-limiting, low ATP concentration (39-49°) ^7^ in which the trailing heads were largely devoid of ADP. This indicates that release of ADP from the trailing head has little effect on the overall geometry of the doubly-attached myosin-5a, in contrast to the small lever swing found for single myosin-5a heads attached to actin when ADP is released ^15, 41^.

The leading heads show an abrupt angle between motor and lever and the lever emerges from the trailing side of the motor, indicating that the converter is in a near-primed position, despite the motor having released phosphate ^7,48^ (Figure 3D). Classification of the leading lever region shows variation in shape, with some levers convexly curved away from the actin filament and emerging from the motor at 120°, rather than straight and emerging from the motor 140° (Figure 3D, right panels). The curved levers still emerge from the trailing side of the motor domain, not the leading side. Thus in these cryoEM data, the leading motor domains are in a near-primed conformation, not unprimed.

The cryoEM data demonstrate that the observations made previously using nsEM ^6,7^ are not artefacts of staining and drying. In particular the two motor domains of the doubly-attached molecule are in different structural states despite being in the same biochemical state. This may underlie their different rates of ADP release.

### Myosin-5a step size distributions from electron microscopy

Myosin-5a molecules with both heads attached on the actin filaments were identified and the separation between the two motor domains of the myosin molecule was measured by an unbiased method, whereby the positions of both motors were determined independently following alignment and classification. For this analysis it was not necessary to determine the polarity of the F-actin filament or to identify which of the two motors was the lead or trail head.

In the cryoEM dataset, motors are separated by 11, 13 and 15 actin subunits (Figure 3A). 77% of molecules have motors attached 13-subunits apart, 10% have motors 11-subunits apart and 13% are 15-subunits apart (Figure 3E). The mean separation is 13.115 actin subunits (= 35.87 nm) (Table 1). A new dataset was also obtained using nsEM of the same myosin-5a construct as used for iSCAT and walking along phalloidin-stabilized F-actin under the same conditions as the iSCAT assay, including 10 µM ATP rather than the 1 µM ATP used in earlier studies ^6,31^. This higher ATP concentration should produce a higher fraction of doubly-attached molecules that have ADP in the trailing motor domain (~50% at 10 µM ATP cf ~10% at 1 µM ATP). Analysis shows 9, 11, 13, 15 and 17-subunit separations (Figure 3C and F). Unlike in longer steps, the global image average of molecules stepping 11-subunits shows the leading head lever emerging from the leading side of the motor domain. This indicates that the smaller separation of motor domains allows the leading motor domain converter to move to the unprimed position, as found previously ^31^ with concomitant distortion in the proximal lever. Like in the cryoEM data, 13-subunit steps are strongly favored (75%), with a mean step length of 13.044 actin subunits (35.87 nm) (Table 1). Thus phalloidin stabilization, which by increasing the rotation per actin subunit ^49^ moves the 13^th^ subunit further from straight ahead, and brings the 15^th^ subunit closer to straight ahead, has little effect on the step-length probabilities.

The EM data complement the iSCAT data by showing the step lengths directly whereas the iSCAT data show strides, which each contain a pair of steps. Both EM methods show a reduced proportion of 15-subunit steps compared with iSCAT.

### How can a straight-walking myosin molecule be taking variable length strides?

If F-actin was a perfectly regular helix, i.e., had a fixed rotation per subunit, our observation of randomly variable step length would imply that the myosin-5a molecule walks drunkenly along actin, moving left and right around the filament as well as along it (Figure 4A). However, in our iSCAT assay, myosin-5a walks straight along actin, staying perpendicular to the plane of the microscope coverslip to which the actin is bound (as described in Andrecka et al.; see also Methods Details). A possible solution to this paradox is that CAD in F-actin could force myosin-5a to vary its step length to continue walking straight. This would be equivalent to someone walking across a river using steppingstones that are variably spaced (Figure 4B). Thus, the relative frequencies of stride lengths would reflect the characteristics of CAD in the F-actin used in the iSCAT assay. We have therefore tested whether CAD in F-actin could be of sufficient magnitude to account for the observed striding behavior. In doing this we have enlarged upon two methods for assessing CAD from images of F-actin ^26^, using recently published cryoEM images of F-actin ^19^ (see Supplementary Information and Figures S4 to S6).

**Figure 4.**
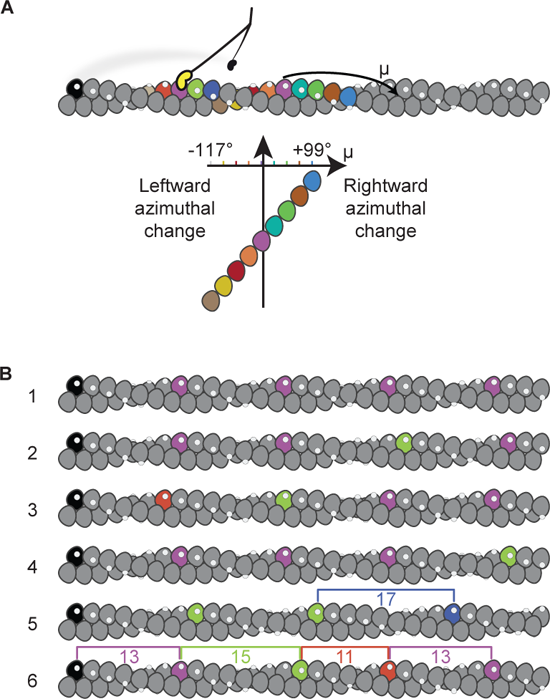
Implications of variable step length if F-actin has fixed or disordered helical parameters. (A) Schematic detailing how varying step and stride lengths would rotate myosin-5a trajectory around the actin filament given a fixed actin rotation per subunit, φ= *−*166.39°. Motor binding site on actin subunit depicted as white spot. A myosin-5a molecule is schematically shown at a stage where one motor (black) has detached from the black actin subunit and the attached motor (yellow) has undergone its power stroke. The detached motor is transiently dwelling off-axis, behind the plane of the figure and will bind to the right of the attached motor. Stride color scheme matches Figure 2. (B) F-actin with variable CAD. Filament 1 has fixed subunit rotation −166.39° and axial translation 2.75 nm. Filaments 2-6 were built incrementally from left to right by adding subunits with an addition to the rotation per subunit randomly drawn from a Gaussian distribution with mean 0° and standard deviation 5.28°. Actin subunits that lie closest to the same azimuth as the zeroth (black) subunit are highlighted with the step colors used in (A), to show expected myosin step lengths, indicated by brackets on filaments 5 and 6.

### Can CAD in F-actin account for the varying stride of myosin-5a?

The implication of CAD for myosin-5a walking is as follows. The mean azimuth (µ_n_) of the n^th^ actin subunit, in degrees, relative to that of a starting subunit having an azimuth (µ_0_) of 0° is given by µ*_n_* = *n*φ modulo 360, where φ is the mean rotation per subunit in the filament. The standard deviation (σ_n_) of that azimuth depends on the RMSD disorder value, *d*, and the number of subunits through *σ_n_* = *d*√*n*. Thus, for any pair of values of φ and *d*, one can estimate the probabilities of the 9, 11, 13, 15 and 17-subunits being closest to an azimuth of 0° and thus the preferred target for straight walking (Figure 5A). It is important to appreciate that on the rare occasions when the 17^th^ subunit is optimally positioned, the 13^th^ will be at µ 55°, requiring a significant reach around the actin. We now apply these principles to test if CAD can account for the relative frequencies of each step and stride length.

**Figure 5.**
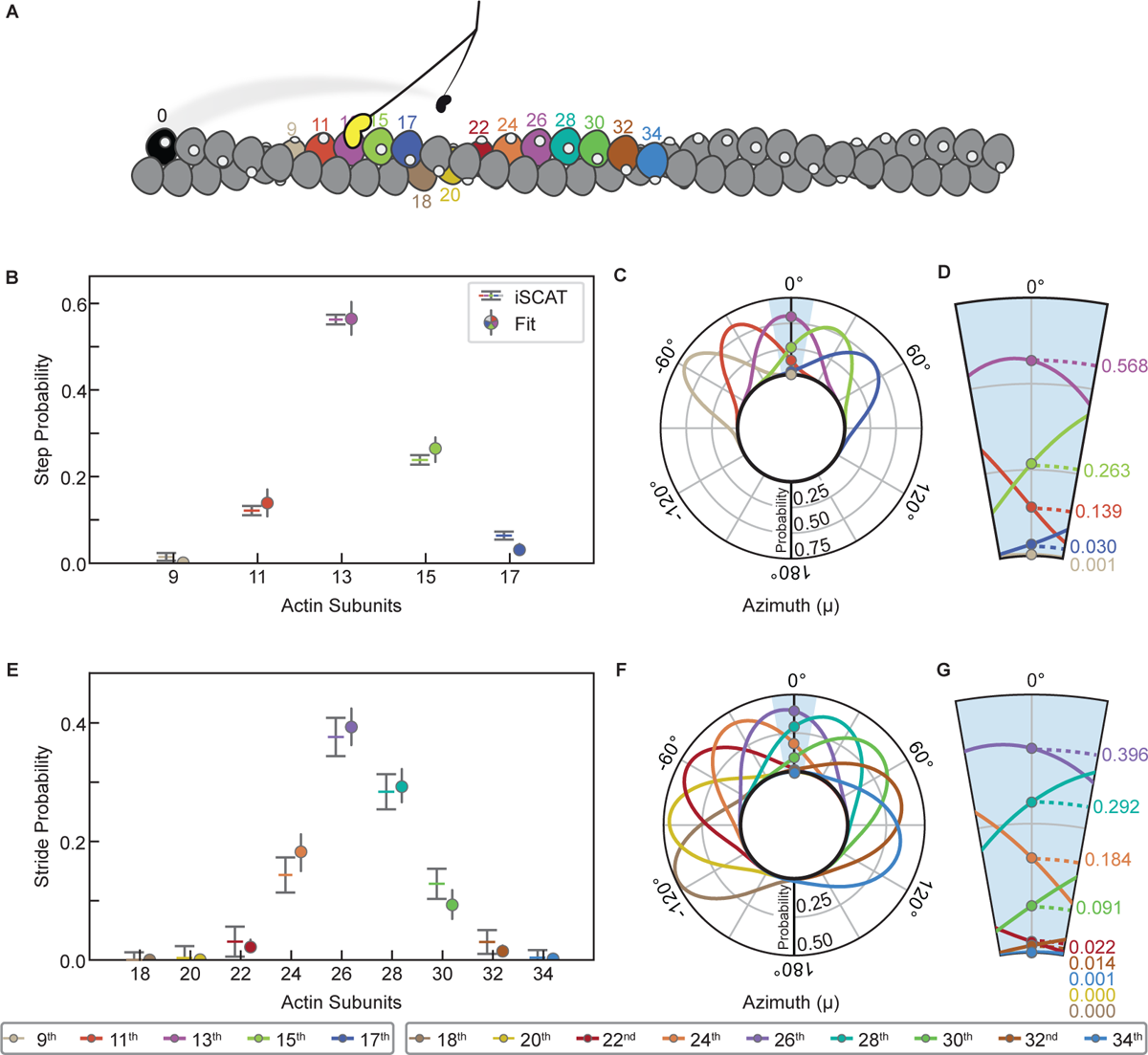
Comparison of F-actin cumulative angular disorder with myosin-5a step and stride length distributions. (A) Schematic numbering the actin subunits involved in steps and strides. Color scheme matches Figure 4. (B) and (E) Calculated probabilities of each step or stride length from iSCAT data, compared with probabilities from the angular disorder 0° azimuth fit, as shown in (C,D) and (F,G), respectively. (C) and (F) Radial plots, i.e., as if looking along the F-actin axis towards the barbed end, showing the azimuthal probability distributions of the n^th^ subunit away from the initial (i.e., the black) subunit along F-actin with the subunit rotation and CAD values (φ = 166.39 5.28°) obtained from least squares fitting to the iSCAT data. Probability densities have been normalized such that the sum of probabilities at µ = 0° is 1. (D) and (G) Expanded ± 10° region showing probabilities of the n^th^ subunit lying at µ = 0°.

If the variable step lengths in the iSCAT data were solely a response to CAD in actin, then our estimates of the relative frequencies of 9, 11, 13, 15 and 17-subunit steps should closely match the relative probabilities of those subunits being correctly positioned for straight walking due to CAD. We therefore used our five stepping frequencies to constrain least squares best fit estimates of φ and *d* in the set of 5 pairs of equations for µ_n_ and σ_n_ to test whether the resulting values agree with current estimates. A robust fit was found, yielding φ= −166.39° 0.11° and *d* = 5.28° 0.26°. Both these values are close to previous estimates, and there is an excellent match between the measured and fitted probabilities of the five step lengths (Figure 5B). Radial plots of the probability distribution illustrate the range of azimuth that each target actin subunit occupies, as viewed along the actin filament axis (Figure 5C). The enlarged segment around µ = 0° shows the azimuthal dependence of probability for straight walking (Figure 5D). For the 17^th^ subunit, the shallow slope of this dependence accounts for the relatively narrow error limits. We conclude that the presence of CAD in F-actin is sufficient to quantitatively account for the variable step lengths taken by myosin-5a.

If CAD is indeed the origin of variable step lengths in the iSCAT assay, then it should also account for the variable stride lengths. The φ and *d* parameters of the actin filament derived from stepping probabilities were therefore used to predict the relative probabilities of the 22^nd^ to 34^th^ subunit positions that are the target sites for the striding motor. These also give a very good match with the observed frequencies (Figure 5E). The radial plots of probability distribution (Figure 5F and G) are noticeably broader than those for the steps, demonstrating the progressive reduction of the correlation with the starting azimuth that is a feature of cumulative (liquid-like) disorder. This good fit to the striding data indicates that CAD in actin does indeed account fully for the spread and relative frequencies of strides observed in the iSCAT data.

Our analysis demonstrates that typical amounts of CAD in F-actin are sufficient to dictate a widely varying stepping pattern for a myosin-5a molecule walking straight.

### Step lengths found by EM indicate myosin-5a walks left-handed around free F-actin

The EM data give a complementary view of myosin-5a stepping to that of iSCAT, in that the myosin is walking along F-actin that is free in solution, rather than apposed to a surface, at the point of EM grid preparation. For the cryoEM dataset, longer steps are rarer than in iSCAT, indicating that myosin-5a prefers to take 13-subunit steps even when CAD means that it must move left-handed around the F-actin axis to do so. The average step length (13.115 actin subunits) is therefore shorter than the average iSCAT step length (13.469 subunits). As a result, using the value of −166.39° for mean actin rotation per subunit, derived above from the iSCAT data, yields an average azimuthal movement that is left-handed of 2.42° per step. This value implies that on average, myosin-5a would take 149 steps to complete a left-handed rotation around the filament, while moving 5.4 µm along it. The nsEM dataset is very similar, yielding an average step length of 13.044 subunits and corresponding values of 2.34°, 154 steps and 5.5 µm.

### Does step length influence myosin-5a ATPase kinetics?

When myosin-5a takes a long step, the levers necessarily lie at a smaller angle to the actin filament than when it takes a short step, as is indeed seen by EM (Figure 3). Since release of ADP from the head moves the lever to a smaller angle to actin ^15, 50^ it might be expected that when the motors are separated by more actin subunits, the rate of ADP release from the trail head would be accelerated, resulting in a shorter dwell time, than would be observed with shorter separations. Conversely, short steps suggest a larger lever angle, slower ADP release and longer dwell time. Because iSCAT resolves varying stride lengths, we have been able to test this idea. There is the complication that a stride may comprise a short step plus a long step in either order, but the shortest strides will comprise only short steps or longest strides only long steps. The two dwell times that refer to a given stride are those of the B state that precedes it and the A state that follows it (see Figure 1E and 6A). Therefore, we have analyzed these dwell times for each stride length.

**Figure 6.**
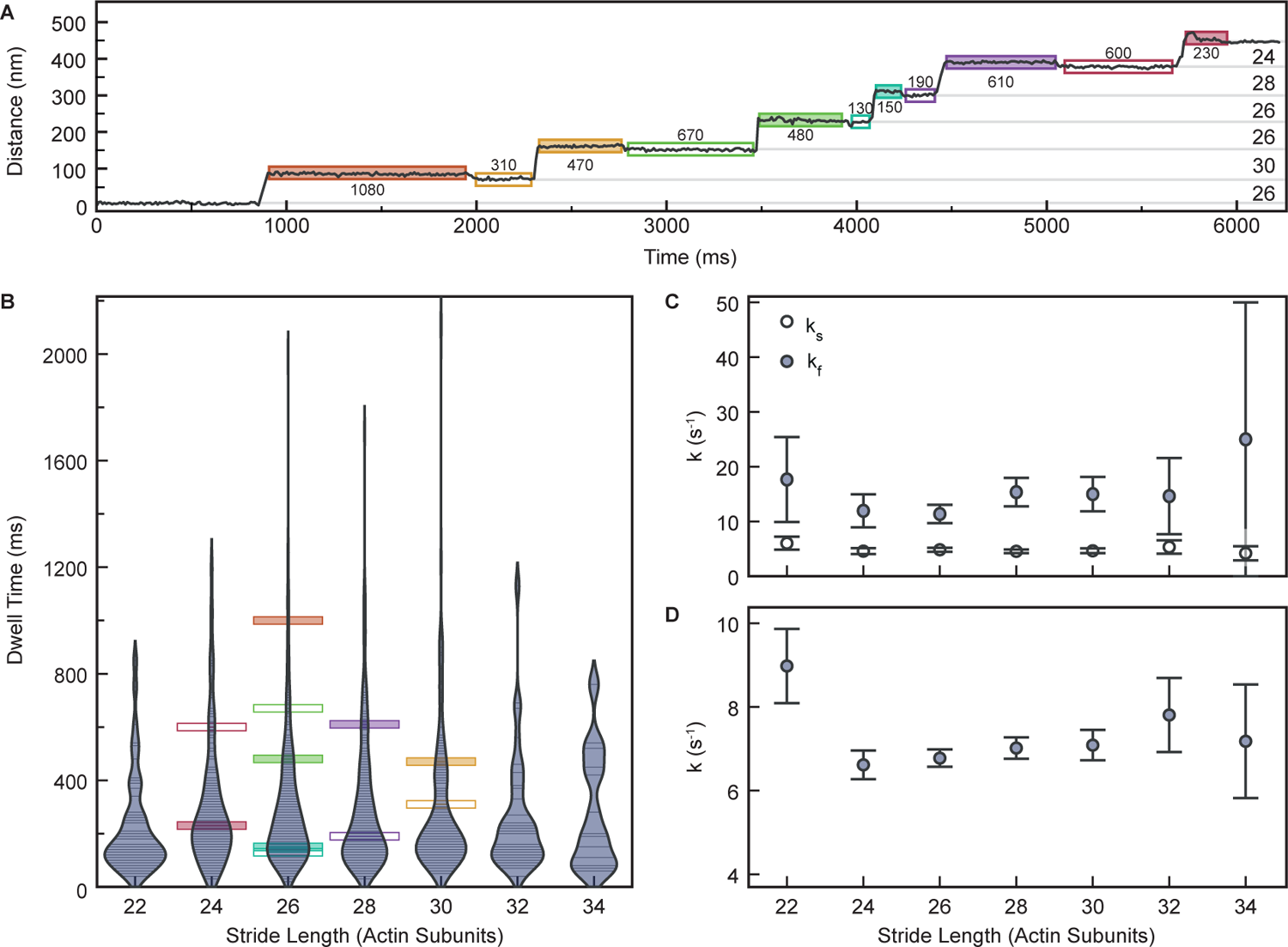
Analysis of impact of stride length on dwell times of A and B states. (A) Distance vs. Time trace of a typical gold-labelled myosin-5a motor domain demonstrating variability in dwell times of labelled motor domain. Dwell times (ms) marked below A states (filled boxes) and above B states (empty boxes). (B) Measured dwell time of A states after stride and B states before stride as a function of stride length. Each observed dwell is shown as a horizontal line. Violin plot envelopes are kernel density estimates of the distributions with bandwidth of 150 ms. (C) Fitted variable rate constants, *k_s_* and *k_f_*, as a function of stride length. (D) Shared fit variable rate constant, *k*, as a function of stride length. Error bars in (C) and (D) denote the standard error of the fit parameters obtained from the observed Fisher information matrix.

We found that there was no significant difference between the dwell times for the A states compared to those of the B states, regardless of stride length, in agreement with our earlier study ^17^, and thus confirming that the gold bead did not affect stepping kinetics. Violin plots of combined A-state and B-state data were similar across all stride lengths, with the most common stride lengths showing the greatest range of values, as expected (Figure 6B). Consequently, there was meagre evidence of dependence of rate constant on stride length (Figure 6C and D, and Figure S7), with rate constants for the two fitting models of *k_s_* ≈ 4.5 s*^−^*^1^ and *k_f_* ≈ 12 s*^−^*^1^ (Figure 6C and S7A) or *k* ≈ 7*s^−^*^1^ (Figure 6D and S7B) and no significant difference between the models as a fit to the data. We conclude that the walking rate, and thus ATPase kinetics, of myosin-5a is not greatly dependent on step length.

## Discussion

Using iSCAT microscopy we have succeeded in resolving a family of stride lengths taken along F-actin by myosin-5a that has one head labelled with a gold nanoparticle. The overall mean and standard deviation of the dataset are similar to those of earlier studies that did not resolve the component peaks, indicating that these variable strides are a constant feature of myosin-5a stepping.

Why were these variable stride lengths not resolved in previous studies? One seemingly trivial reason is that because the strides differ in length by one actin subunit distance (5.5 nm) it is necessary to aggregate the data into small enough bins (≤1.8 nm) to resolve that spacing, yet this has generally not been done. A second reason is that the method used to determine the start and end positions of each stride can add sufficient error that the peaks overlap. Thus, for the iSCAT data, a switch from measuring lengths from distance-time traces ^17^ to measuring them from the x, y coordinates proved critical to resolving the strides. A further challenge arises where the center of mass of the molecule is being monitored, such as if both heads are labelled or in optical trap studies. Only if successive motor separations along actin are all the same is the movement of the molecule equal to the motor separation. If they differ, as we have shown they do, then in making the transition between them, the center of mass moves by the average of the two separations. For example, if a motor separation of 13-subunits is followed by one of 15-subunits, the center of mass moves 14-subunits. This means that the movement of the center of mass is not directly reporting the sizes of the steps taken by the motors. It also means that it is more demanding to resolve the subpeaks because they are only 2.75 nm apart.

### Misconception of the myosin-5a average step length explained

In contrast to common statements that myosin-5a walks by taking 36 nm steps along actin, we find that only about half the steps span the canonical 13-subunits (35.75 nm). Almost a quarter span 15-subunits with progressively smaller proportions spanning 11, 17 and 9-subunits. Because of this diversity of step lengths, neither the average step nor average stride length corresponds to an actual movement, as these average values fall between the separation of actin binding sites. These results from the iSCAT assay pertain also to previous assays that used F-actin attached to a coverslip or suspended between a pair of beads in the optical trap. This is because in each case only a limited azimuth of F-actin is available for the myosin to walk along. The presence of a family of unresolved steps or strides in the previous data explains why the breadth of the observed distribution was broader than would have been expected from the resolution of the methods used.

### Cumulative angular disorder of actin filaments accounts for myosin-5a step and stride frequency

We show that CAD exists within specimens of F-actin prepared by modern methods. Remarkably, CAD is of sufficient magnitude to account for the relative frequencies of the step and stride lengths found in the iSCAT data for myosin walking along a single azimuth of the actin filament. This nicely resolves the paradox of how myosin-5a could take steps of varying length while still walking straight on the helical F-actin. Recent high resolution cryoEM structures of F-actin ^19, 20^ omit this feature of F-actin structure, instead emphasizing a precise (average) value for subunit rotation (close to −166.5° per subunit). However to obtain high resolution, these studies restrict the reconstructed volume to a few subunits which reduces the impact of disorder, and when a longer segment of filament is used, resolution reduces ^19^, as is expected from CAD.

Our study has revealed the biological significance of CAD in influencing the behavior of myosin motors. Although CAD has been well-characterized for F-actin purified from skeletal muscle ^26, 27^, there are no data on CAD in cytoskeletal actin isoforms ^51^, and also no data for the influence of cytoskeletal tropomyosin or for CAD in actin filaments bundled by cross-linking proteins such as fascin or fimbrin. Although for some of these complexes there are data on filament flexibility, there is no established causal linkage between the magnitude of CAD and flexibility. Our study therefore indicates a need for further research to characterize CAD in the cellular environment and to understand the roles of CAD in determining cellular behavior. Understanding the temperature dependence of CAD magnitude and dynamics would also be beneficial for understanding the importance of CAD in F-actin under physiological conditions for warm- and cold-blooded animals.

### Myosin-5a walks left-handed around F-actin

Previous studies have directly shown that myosin-5a molecules can walk left-handed around a suspended actin filament that has been stabilized by phalloidin ^52–54^. Phalloidin increases actin subunit rotation to −167° ^49^. This moves the 13^th^ subunit to an average azimuth 11° left of straight ahead and correspondingly the 15^th^ subunit moves to be only 15° to the right. Thus, walking straight on such filaments requires a high proportion of 15 subunit steps. For myosin taking mainly 13 subunit steps, as we observe by both cryoEM and nsEM, the molecule would twirl left-handed around the filament more strongly than for phalloidin-free F-actin. This may account for the shorter pitch of twirling (2-3 µm) ^52–54^ compared to our prediction of 5.5 µm. This can be compared to an average run length of ≈1.3 µm ^35, 55, 56^, from previous studies. The earlier conclusion that left-handed twirling along phalloidin-stabilized F-actin implied that myosin took 11 and 13 subunit steps ^52^ is incorrect, because it assumed −166.154° rotation per subunit for F-actin (i.e. 13/6 helical symmetry) rather than −167°.

### Structure of the walking myosin-5a molecule

It is perhaps surprising that this is the first report of a cryoEM study of myosin-5a walking on actin, but in practice we have found it difficult to get the walking molecules into the holes of the EM support film. Nevertheless, our small dataset clearly shows that the converter of the leading motor domain is held in a stressed, near-primed position through tethering to the trail head. Thus, the flash-frozen, unstained walking molecule appears similar to our previous studies using nsEM at very low ATP concentrations ^6,31^, and also to the nsEM data presented here that were obtained at ten times higher ATP concentration than previously.

The ‘Telemark skier’ analogy used to describe the appearance of doubly-attached myosin-5a ^6^ has since been interpreted by some to mean that both motor domains are in an unprimed conformation, with the lead lever bent back to create the knee of the skier ^33^. The EM data presented here confirms that this is not the case. The leading lever is not bent. Instead, the knee of the leading leg of the skier is actually the lever-motor junction, with the lower leg being formed from the motor domain together with the actin subunit it is attached to and appearing angled backwards because the motor is attached to the leading side of the prominent subdomain 1 of actin. In contrast, in the trailing head, actin subdomain 1, the motor domain and the lever are all roughly colinear, creating the different shape resembling the trailing leg of the skier. As the powerstroke of myosin-5a is similar in distance to the canonical step length of 35.75 nm ^57^, a primed myosin head could easily bind to create the next leading head, with the subsequent rapid release of phosphate ^48^ generating stress within the doubly-attached molecule as the head is unable to adopt the preferred unprimed (post-powerstroke) structure of actomyosin.ADP.

Our nsEM, performed at the same ATP concentration as the iSCAT assay shows that for 11-subunit motor separation this tethering is insufficient to prevent converter movement to the unprimed position, with concomitant distortion of the leading lever but little change in the trailing lever (Figure 3C). This may explain why we find that the stepping rate is not slower for short steps, but it is unknown whether this also reduces gating of lead motor ADP release.

### Mechanical properties of myosin and impact on ATPase kinetics

Our analysis indicates that myosin-5a has to take variable length steps in the iSCAT assay because the sample geometry restricts azimuthal variation, forcing the motor domain to attach to the subunit of the disordered actin filament that is closest to straight ahead. Thus, the mechanical properties of the myosin cannot be directly assessed, only that there is sufficient compliance within the molecule to allow it to vary its step length over the range observed. However, a similar range of step lengths, albeit that 13-subunit steps are more strongly favored, is found by both cryoEM and nsEM, in which the myosin is free to explore around the actin filament to locate binding sites. This similarity indicates that myosin-5a has a preferred lever-lever angle that places the motors ≈36 nm apart, as we previously proposed from iSCAT data ^17^. The greater frequency of 13-subunit steps in the EM data, unaffected by phalloidin stabilisation of F-actin, indicates that when the myosin is free to explore around the actin filament to overcome the effects of CAD, there is a still a preference for binding with that motor separation, reinforcing the idea of a preferred geometry for the singly-attached molecule.

The symmetrical Gaussian spread of the frequencies of shorter and longer steps further suggests a thermally-driven fluctuation in motor separation that allows both longer and shorter steps. This could arise from bending within the levers and at the lever-motor junctions and from fluctuations in the lever-lever angle. The standard deviation of the distribution of step lengths leads to an estimate for the stiffness of the molecule along the actin filament axis, through the Equipartition Principle, as stiffness, *κ* = *k_B_T/σ*^2^. For our nsEM data, σ= 3.289 nm (Table 1) yielding *κ* = 0.374 pN/nm. If we suppose that the lever-lever angle is fixed, then since the two heads are mechanically in series, the stiffness of each head would be 0.748 pN/nm. A similar standard deviation (3.0 nm) was reported by Oke ^58^ also using nsEM, indicating a similar stiffness.

Because we have been able to examine the dependence of stepping kinetics on stride length, we can use this estimate of myosin-5a stiffness to compare our data with earlier optical trap measurements of the impact of external force on kinetics of ADP release and ATP binding by single myosin-5a heads ^15, 59, 60^. Our EM data indicate that a 13-subunit motor separation has least strain. A 15-subunit separation implies a forward displacement of the lever-lever junction of 2.75 nm, and thus an assisting force on the trailing head of 2.75 x 0.374 = 1.03 pN. For a 17-subunit separation the force would be 2.06 pN, which would be expected to produce a marked acceleration of ADP release and thus reduction in B-state dwell time. It remains to be understood why this acceleration was not detected, but we note that in the iSCAT assay the orientation of the head with respect to the actin filament axis is closely specified (and relevant to myosin walking in the cell), whereas in the optical trap the orientation of the head is not known.

## Conclusions

Our demonstration of variable step and stride lengths for myosin-5a provides a framework for understanding how the molecule manages to transport cargoes through the complex and crowded cytoskeletal matrix. The steps the molecule takes are constrained by the mechanical properties of the myosin and, as we have revealed, also by CAD in F-actin. Unlike in macroscopic transport systems, both motor and track are ‘soft’, containing elements of random disorder that create adaptability.

## AUTHOR CONTRIBUTIONS

**Conceptualization**: Y.T., J.A., J.R.S., P.K. and P.J.K.; **Methodology**: A.F., Y.T., K.T., J.A., S.A.B., J.R.S., P.K. and P.J.K.; **Software**: A.F., D.C., N.B., S.A.B. and G.Y; **Validation**: A.F., Y.T., K.T., J.A. and N.B.; **Formal analysis**: A.F., Y.T., K.T., J.A., N.B., G.Y., S.A.B., A.P.C. and P.J.K.; **Investigation**: A.F., Y.T., K.T., J.A. and N.B.; **Resources**: A.F., Y.T., K.T., J.A., D.C., G.Y., J.A.H., J.R.S., P.K. and P.J.K.; **Data curation**: A.F., J.A., N.B. and P.J.K.; **Writing – Original Draft**: Y.T., J.A. and P.J.K.; **Writing – Review and Editing**: All authors; **Visualization**: A.F., J.A., N.B. and P.J.K.; **Supervision**: Y.T., J.A., J.R.S., P.K. and P.J.K.; **Project Administration**: Y.T., P.K. and P.J.K.; **Funding Acquisition**: J.R.S., P.K. and P.J.K.

## ACKNOWLEDGEMENTS

We thank the Electron Microscopy Facility of the National Heart, Lung, and Blood institute (NHLBI) for the use of facility. We thank Fang Zhang for her help with reagent preparation. We thank Vitold Galkin for clarifying the properties of CAD and Greg Alushin for encouragement to look for CAD in his archived cryoEM F-actin data. This work was supported by the following funding: Biotechnology and Biological Sciences Research Council (Grant BB/S015787/1 to A.P.C.); Intramural Research Program of the NHLBI, NIH (ZIA HL004229 to J.R.S.); Wellcome Trust (Grant 076057 to P.J.K.); European Research Council (ERC) Consolidator Grant (PHOTOMASS 819593 to P.K.), and the Engineering and Physical Research Council (EPSRC) Leadership Fellowship (EP/T03419X/1 to P.K.). For the purpose of Open Access, the author has applied a CC BY public copyright license to any Author Accepted Manuscript (AAM) version arising from this submission.

## COMPETING FINANCIAL INTERESTS

The authors declare no competing interests.

## Supplementary Information

### Characterizing cumulative angular disorder in F-actin

CAD has not been mentioned in recent high-resolution studies of F-actin structure ^1–4^, we therefore first tested whether it could also be detected in F-actin prepared and imaged using current cryoEM methods. We analyzed recent high resolution cryoEM micrographs ^1^ from the EMPIAR archive, developing two tests for CAD ^5^ that make use of the spacings between crossovers of the two long-pitch helices of actin subunits. To properly interpret the results, we needed also to understand the impact on these tests of observer error in picking the position of crossovers in F-actin images, as this had not previously been evaluated. We therefore also analyzed test data from model actin filaments that were created in silico to have specified magnitudes of CAD (Figure 4B) and position error from picking crossovers (see Methods Details). Since both these variables were assumed to have a Gaussian spread of values distributed randomly along the filament, these simulations were done using large numbers of model filaments and the results averaged, in order to reduce the impact of the random fluctuations between filaments (Figure S5 and S6).

### Test 1: Consecutive crossover correlation

The first test examines the relationship between successive crossover spacings by creating a scatter plot from pairs of adjacent crossover spacings (Figure S4B). For a perfect helix, measured without error, all spacings will be identical (Figure S4B i). Introducing position error in the measurement, which generates a crossover spacing error (see Methods Details) along a perfect helix, the data points cluster into an ellipse with major axis having slope −1, because an overlong crossover estimate tends to be followed by a short one (and vice versa) (Figure S4C ii). The result is that the standard deviation of the data projected onto an axis of slope −1 (SD_-1_) is much greater than that projected onto the orthogonal axis of slope +1 (SD_+1_). The ratio of the two SDs (SD_-1_/ SD_+1_) is 1.73 (Figure S4C ii, inset), and is essentially independent of the magnitude of error from picking (Figure S5B). In contrast, the randomness inherent in CAD scatters the points in all directions from the mean center (Figure S4B iii). The non-linear relationship between crossover spacing and subunit rotation angle results in a SD slightly higher on the axis of slope +1 than on the orthogonal axis of slope −1, yielding a SD ratio of 0.97 (Figure S4C iii, inset). This value is, again, invariant with CAD magnitude (Figure S5D). As expected, a combination of CAD and position error produces an intermediate SD ratio between the two limiting values (Figure S4C iv). This modelling thus shows that an SD ratio below 1.73 is an indicator of CAD, but does not directly yield a value for CAD because the magnitudes of CAD and position error interact to determine the net SD ratio (Figure S5B and D).

In the cryoEM images, the polarity of each filament could be assigned by eye and the locations of the series of crossovers were clear (Figure S4D). For n = 371 crossovers picked in n = 54 actin <u>fil</u>aments, the mean spacing (^-^x) was 37.95 nm with an SD of 2.19 nm. The mean spacing implies a mean rotation per subunit (φ) of −166.96°, close to the value (−166.85°) reported for these images ^6^. The scatter plot of successive crossover spacings yielded a SD ratio of 1.36, well below the value for position error (1.73), and this thus indicates a strong contribution from CAD (Figure S4E).

### Test 2: Cumulative variance

The second test for CAD derives from its cumulative nature. For a family of filaments, the set of measurements of the distance along a filament between crossovers separated by a given number of crossovers will have a mean and SD. For a perfect helix picked with zero error, the SD will always be zero (Figure S4C i). Modelling confirms that when position error is added, the SD of crossover distance does not depend on how many crossovers intervene, because the magnitude of SD derives only from the position errors at the start and end crossover locations in the image (Figure S4C ii). For a helix with CAD there is additional variability because the crossover spacings really are shorter and longer. The further one progresses along the filament, the broader the range of possible crossover positions will be and thus the SD of crossover position rises with crossover number (Figure S4C iii). More detailed analysis of large numbers of long model filaments and using a range of CAD magnitudes confirms the rise in SD and also shows that the variance of crossover distance rises linearly with crossover number (Figure S6A and B). The SD of crossover distance rises linearly with CAD over the range 0 – 10° (Figure S5F). Position error increases the SD through an addition to the variance value. It thus adds a constant offset to the variance values and has no effect on the slope (Figure S6C). Helical geometrical arguments relate crossover spacing to subunit rotation and this in turn allows a least squares estimation of both CAD and position error (see Methods Details).

Applying this test to the cryoEM data, the SD of crossover distance does indeed increase with the number of crossovers traversed, confirming that CAD is present in the F-actin (Figure S4F). The data points are noisy because of the random fluctuations expected within the relatively small dataset (n=54 filaments analyzed). The least squares fit yielded an estimate of 2.24° per subunit for CAD, with a crossover error SD of 1.30 nm, implying a position error of 0.92 nm (Figure S5F). Applying these estimates to the first test (above) predicts a SD ratio of 1.31, close to the value of 1.36 observed.

Therefore, we conclude that CAD is an overlooked feature of F-actin. The SD of crossover spacing in this cryoEM dataset (2.19 nm) is smaller than in an early study (5.4 nm ^7^), indicating that CAD magnitude is sensitive to experimental conditions.

### Contact for reagent and resource sharing

Further information and requests for reagents and resources should be directed to and will be fulfilled by the lead contact, Yasuharu Takagi.

## KEY RESOURCES TABLE

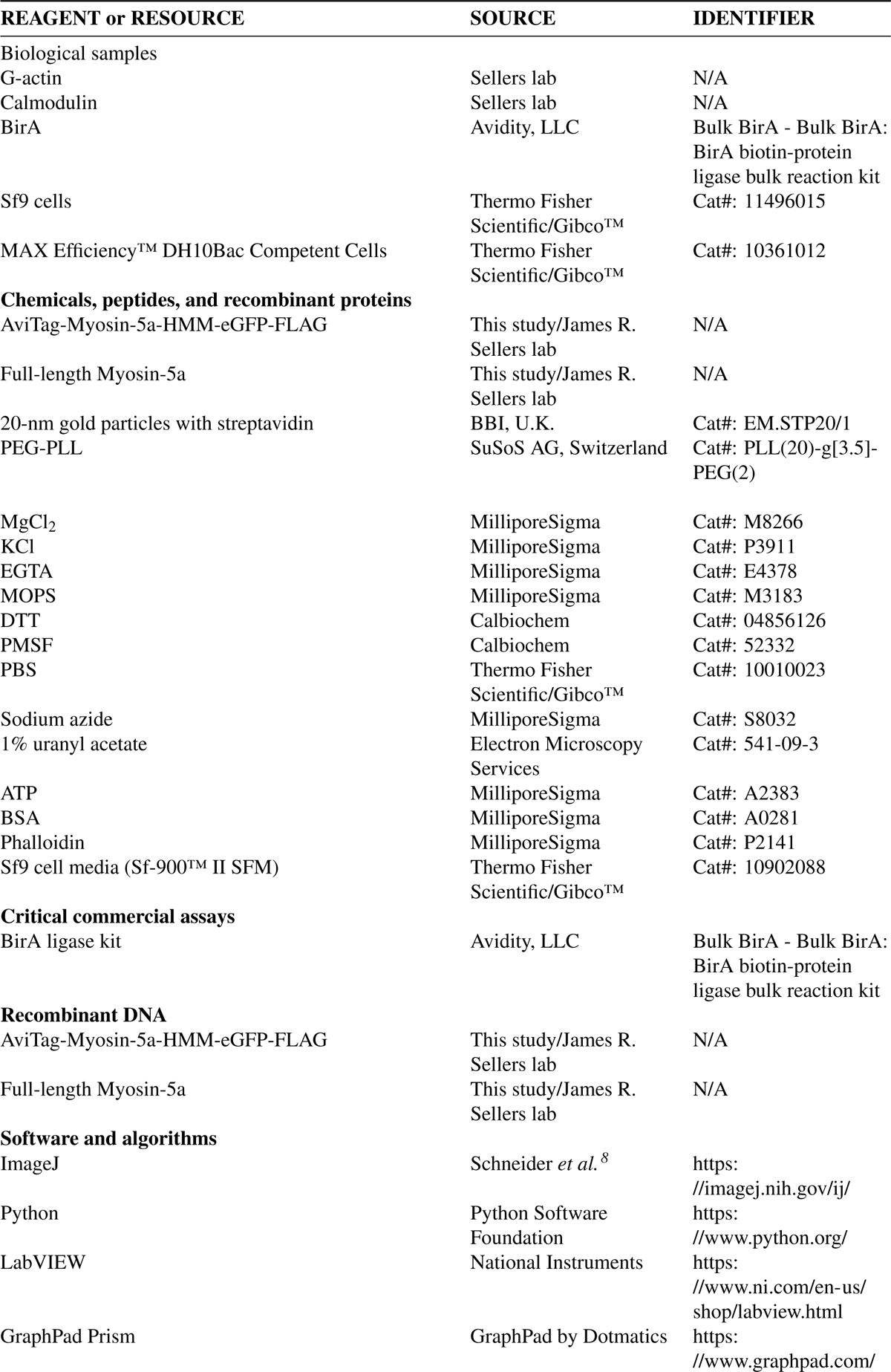
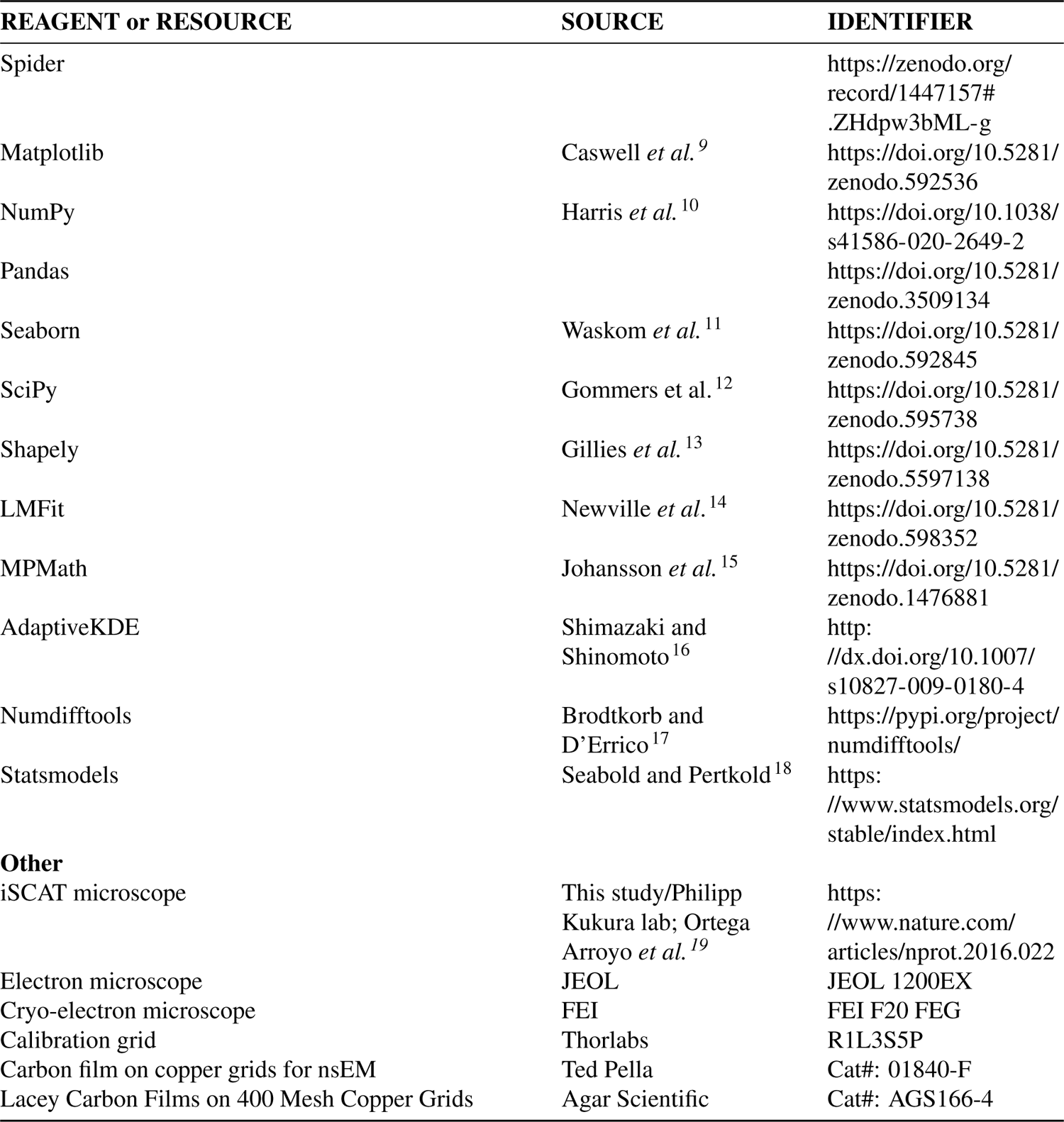

### Experimental model and subject details

Plasmid DNA for recombinant protein was amplified in E. coli DH5a strain (Thermo Fisher Scientific) in LB medium at 37°C overnight. Plasmid DNA was transformed into DH10Bac E. coli cells (Thermo Fisher Scientific) and then used to make baculoviral stock per the manufacturer protocol with minor modifications ^20^. Recombinant proteins were overexpressed using the Sf9/baculovirus expression system using the Sf900 II medium (Thermo Fisher Scientific). The cells were grown in a temperature-controlled incubator shaker at 27°C/90 RPM in 2.8 L Fernbach flasks without baffles for 48 - 60 hrs. Sf9 cells were collected into a 50 ml conical tube and flash frozen with liquid nitrogen. Cell pellets were stored in the −80 °C freezer for long-term storage until purification was performed.

## Methods Details

### Protein purification

Rabbit skeletal muscle G-actin was prepared as described ^21^ and stored in liquid nitrogen until use. For cryoEM and nsEM, F-actin was prepared by a two-step procedure in which the divalent cation in the G-actin is first exchanged from Ca^2+^ to Mg^2+^ by addition of 0.1 volume of exchange buffer (3 mM MgCl_2_, 11 mM EGTA, pH 7.0) in ice for 15 minutes, then polymerized by addition of 0.1 volume of polymerization buffer (300 mM KCl, 12 mM MgCl_2_, 1mM EGTA, 120 mM MOPS, pH 7.0), yielding a solution composition of 25 mM KCl, 1.25 mM MgCl_2_, 1.0 mM EGTA, 0.17 mM ATP, 10 mM MOPS, 0.17 mM Tris, pH 7.0.

For the iSCAT measurements and nsEM, a mouse myosin-5a HMM-like construct ((i.e. having the two heads together with the proximal coiled coil, like heavy meromyosin derived from muscle myosin-2; myosin-5a sequence amino acid sequence 1-1090) with an N-terminal AviTag biotinylation sequence ^22, 23^ and C-terminal eGFP and FLAG sequence was expressed in the presence of calmodulin, using the Sf9/baculovirus expression system and purified as described previously ^24^. The following is the amino acid sequence of the construct (total: 1352 amino acids):

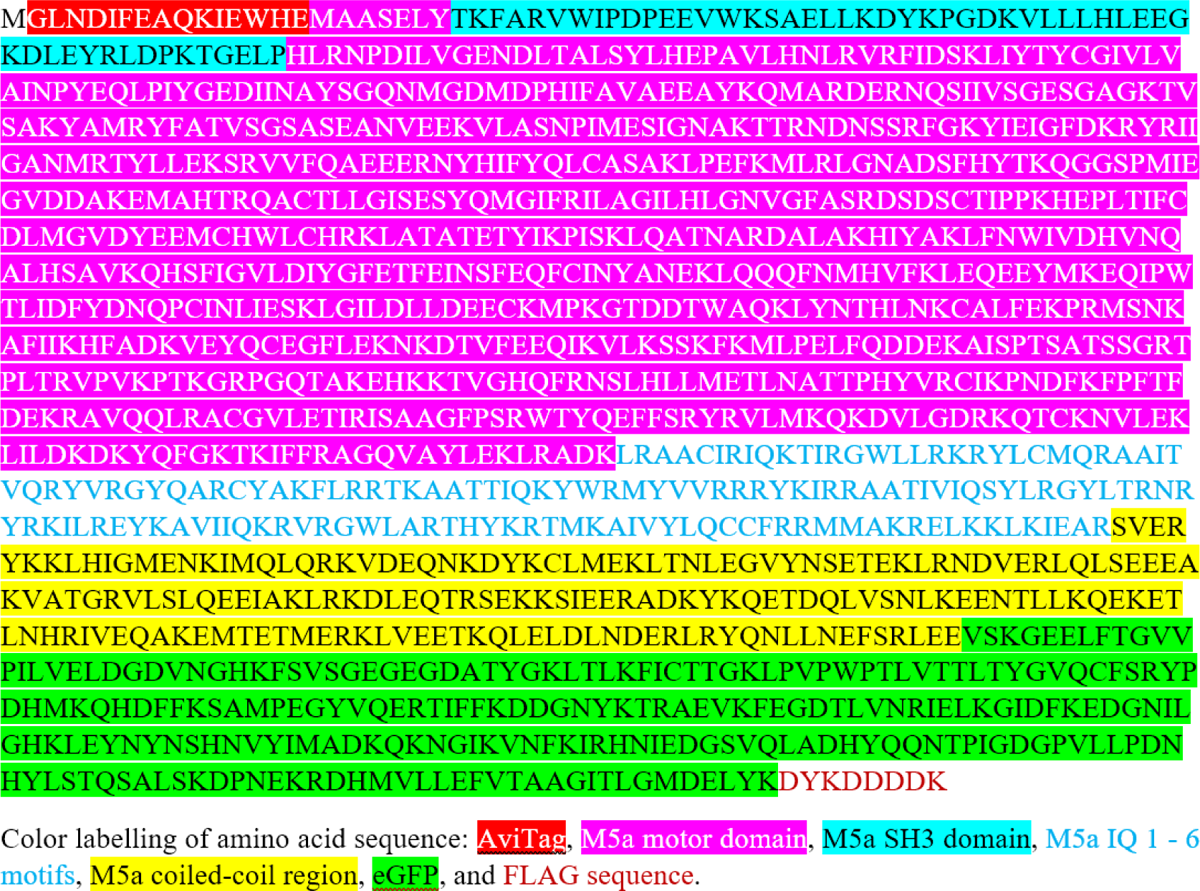

For cryoEM, the melanocyte-specific isoform of full-length mouse myosin-5a heavy chain ^25^ was expressed together with calmodulin in Sf9 cells, purified using an N-terminal FLAG tag, dialyzed against 100 mM KCl, 0.1 mM EGTA, 1 mM DTT, 0.1 mM PMSF, 10 mM MOPS, pH7.0 and stored drop-frozen in liquid nitrogen, similar to as described before ^24^. It was incubated with 4 moles of purified calmodulin per mole myosin overnight on ice, prior to cryoEM data collection.

### Single molecule motility assay using iSCAT microscopy

Biotinylation of the myosin-5a HMM construct (~100µ*l* of ~ 4 – 5 µM) with BirA ligase (Avidity, Aurora, Colorado), was performed on ice for ~12 hours as described in detail by the manufacturer protocol (https://www.avidity.com/showpdf.asp?N=B9B6C28E-1976-4CFE-89DE-E37D1D8DB2CA). After this reaction, dialysis against 1 L of buffer containing 50 mM KCl, 0.1 mM EGTA, 1 mM DTT, 10 mM MOPS, pH 7.3, was performed for ~30 minutes (2x; total ~1 hr), to remove excess biotin. Biotinylated myosin-5a HMM was then used immediately for the iSCAT assay, or aliquoted and frozen in liquid nitrogen and stored in either liquid nitrogen storage or the −80 °C freezer for future use.

Newly biotinylated or stored myosin-5a HMM sample (750 nM per molecule) was diluted 10x in motility buffer (MB; 40 mM KCl, 5 mM MgCl_2_, 0.1 mM EGTA, 5 mM DTT, 20 mM MOPS, pH 7.3) supplemented with 0.1 mg/ml BSA and 5 µM calmodulin. This was kept as a stock myosin solution (75 nM), on ice for the day of the experiment.

To form the stock myosin-5a HMM/gold nanoparticle complex, 0.77 nM streptavidin-conjugated 20 nm gold nanoparticles in tris-buffered saline (pH 8.2) containing 10 mg/ml BSA and 15 mM NaN_3_ (BBI, UK) was incubated with 0.25 nM biotinylated myosin-5a HMM in MB. Thus, the gold:myosin ratio was ~1:0.3, similar to previous published methods ^26, 27^ ensuring that on average, one or no myosin-5a molecule attached per gold nanoparticle. Myosin-5a HMM/gold nanoparticle complex was incubated on ice for approximately 15 min, immediately before preparing the final assay mix. New complexes were made prior to preparing a new flow cell for the assay.

20 µM F-actin stock solution was prepared in polymerization buffer (50 mM KCl, 1 mM MgCl_2_, 1 mM EGTA, 10 mM imidazole (pH 7.3), 1.7 mM DTT, 3 mM ATP) and diluted with MB to 0.2 - 1 µM prior to use.

The flow cell (volume ~20 µl) was prepared as described ^28, 29^, first rinsing with 1 mg/ml poly-(ethylene glycol)-poly-L-lysine (PEG-PLL) branch copolymer (SuSoS AG, Switzerland) in PBS (1 mM KH_2_PO_4_, 155 mM NaCl, 3 mM Na_2_HPO_4_-7H_2_O, pH 7.4; Gibco) and incubated for 30 min. The flow cell was washed twice with MB (100 µl per wash) before adding the F-actin.

After F-actin was pipetted into the flow cell, attachment of the F-actin to the surface of the coverslip was monitored *via* the iSCAT microscope, as it can monitor label free proteins ^30^. The chamber was then washed extensively with MB (500 µl) and the surface was blocked additionally with 1 mg/ml BSA in MB (100 µl). Prior to making the final assay mix, 0.1 M ATP (pH 7.0) was diluted 100x in MB to 1 mM ATP. This was then further diluted 100x to 10 µM ATP in the final assay mix.

The final assay composition in which stepping of myosin-5a HMM with a gold nanoparticle attached was observed, was as follows: 77 pM 20 nm gold nanoparticle attached with 25 pM myosin-5a HMM (*i.e.*, 10x dilution of previously mentioned stock), 10 µM ATP, 5 µM calmodulin, 40 mM KCl, 5 mM MgCl_2_, 0.1 mM EGTA, 5 mM DTT, 20 mM MOPS (pH 7.3). Data collection was performed at room temperature (19 – 22 °C) and flow cells were not used for more than 30 minutes.

### Description of iSCAT microscopy and data collection

Interferometric scattering microscopy was performed as described previously ^31, 32^. In iSCAT microscopy, a 445 nm diode laser beam is scanned across the sample using a pair of orthogonal acousto-optic deflectors (Gooch & Housego) mapped into the back focal plane of the objective (PlanApo N 60x, 1.42 NA, Olympus) with a 4f telescope, giving an approximately 30 x 30 µm illumination. The illumination and detection paths are separated using a quarter-wave plate beneath the objective and polarizing beam splitter after the 4f telescope. The detected signal is imaged onto a CMOS camera (MV-D1024-160-CL-8, Photonfocus) at approximately 333× magnification (31.8 nm/pixel). The true magnification was determined using a calibrated resolution grid (Thorlabs) (Figure S1D).

Because of the potential for creating mirror inversions of the iSCAT image, both in the iSCAT microscope and in the analysis software, we confirmed that a clockwise, circular translation of the sample (as viewed from above the sample, *i.e.,* the coverslip on the far side of the actin filaments) resulted in a clockwise rotation of the iSCAT image and that this was unchanged by the software. By combining this information with the fact that myosin-5a always processes towards the barbed end of actin, we define left and right relative to an actin filament as if viewing the filament and motor from above, with the actin filament aligned along the y-axis and myosin walking up the y-axis.

### nsEM sample preparation and data collection

For nsEM, 20 µM F-actin was stabilized with 24 µM phalloidin. F-actin and the myosin-5a HMM construct used for iSCAT assays, but without the gold bead, were each diluted in 50 mM NaCl, 1 mM MgCl_2_, 0.1 mM EGTA, 10 µM ATP, 10 mM MOPS, pH 7.0. Equal volumes of F-actin and myosin were then mixed to give final concentrations of 500 nM and 25 nM respectively. Samples were incubated for 30s, applied to UV-treated carbon film on copper EM grids and stained using 1% uranyl acetate. Micrographs were recorded on a JEOL 1200EX microscope using an AMT XR-60 CCD camera at 0.31 nm/pixel, calibrated using catalase crystals.

### CryoEM sample preparation and data collection

Full-length mouse myosin-5a in 100 mM KCl, 0.1 mM EGTA, 1 mM DTT, 10 mM MOPS, pH7.0 was incubated with 4 moles calmodulin per mole myosin overnight in ice. ATP was added from 7.8 mM stock solution to give 0.2 mM ATP, then mixed with a small volume of 83 µM F-actin and applied within 10 s to a lacey carbon grid (glow discharged in amylamine), quickly blotted with Ca-free paper and flash frozen in ethane slush. Final conditions: 0.83 µM myosin-5a, 3.6 µM calmodulin, 2.1 µM F-actin, 0.20 mM ATP, 95 mM KCl, 48 µM MgCl_2_, 0.12 mM EGTA, 10 mM MOPS, pH 7.0 at 22°C. Specimens were imaged in an FEI F20 FEG cryoEM at ~4 µm underfocus and recorded as 2×2k CCD images at 0.589 nm/pixel, calibrated with catalase crystal.

## Quantification and Statistical Analysis

### iSCAT stride length analysis

iSCAT images were collected at 200 Hz and averaged down by a factor of 2 to identify the AB transitions (Figure 1C). The imaging frame rate was limited to 100 Hz (although iSCAT detection does allow much faster rates) to reduce the effects of label-linker flexibility, which increase localization noise at higher speeds and reduce our ability to identify the AB transition. An estimate of the localization precision achieved under these experimental conditions was determined by the positional fluctuations (σ ~ 0.91 nm, SD) ^26^ of a surface-attached label recorded under the same conditions as the trajectory in Figure 1C. For actin-bound myosin-gold conjugates the localization noise increases (σ = 1.6 ± 0.3 nm) ^26^, but remains on average ~2 nm which allows detection of the AB transitions on the order of 5 nm.

The pixel values of the gold particle were fitted with a 2-dimensional Gaussian to determine its x, y position (Figure 1C and D and Figure S1B and C) at nanometer precision. Myosin-5a strides were measured as the distance between the centers of two successive B states. The start and end of each B state was identified using a step detection algorithm ^33^. The mean x, y coordinates (*x_i_, y_i_*) of each B state and the Euclidian distance between these mean x, y coordinates calculated:

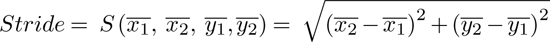

The data can be visualized as either an xy-plot (Figure 1C) or a distance-time trace (Figure 1E). To plot a distance time trace, the x and y values of the first data point in the run, at time = 0, (x_0_, y_0_) were subtracted from each of the values in the run (x, y) followed by addition in quadrature:

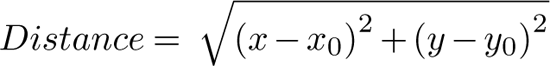

However, distance-time traces do not adequately represent behavior during a dwell period because there is systematic underestimation of the contribution of bead movements orthogonal to the line joining the first data point to the mean center of the dwell position.

Myosin-5a strides, illustrated as blue arrows on the schematic, (Figure 1B) and highlighted as blue transitions on the example data trace (Figure 1C and E) were clear from these trajectories, as were the AB transitions, illustrated as red arrows on the schematic (Figure 1B) and highlighted as red portions on the example data trace, where they show as a decrease in distance (Figure 1C and E), as previously published ^26^. The spread of particle localizations was calculated as the standard deviation of the gold particle during its stationary state (B-state, in this case) (Figure 1C), the uncertainty of particle positions is reported as the standard error of the mean (Figure 1D). The error on stride distances (*σ_Stride_*) (Figure 1D) was calculated by error propagation of the standard errors of the means:

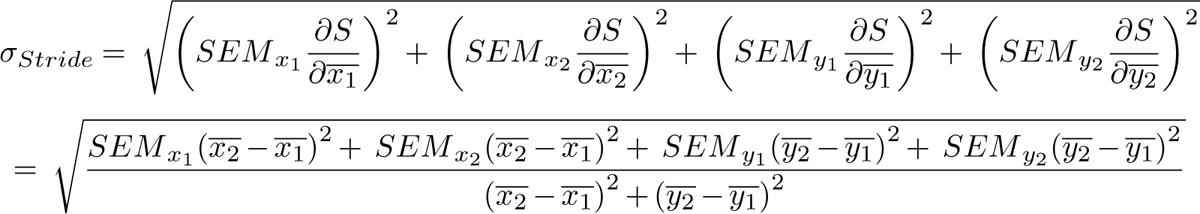

### Orientation of myosin-5a in the iSCAT assay

To fully understand the geometric implications of the experimental results, one must consider the geometry of the myosin-5a molecule relative to the experimental imaging axis. It is possible to rule out the scenario where the myosin-5a molecule binds and remains parallel to the glass surface, along the side of actin filament, for the entirety of its processive run. Due to BSA surface blocking, possible myosin binding sites on the side of the actin filaments are far more inaccessible as compared to those on the top. Binding of myosin-5a on top of the filaments will therefore minimize surface interactions with the gold label. Additionally, through experimental comparison of myosin-5a which has been labelled either at the motor or at the end of the coiled-coil domain, a significant contrast difference is seen between the motor-bound and coiled-coil-bound gold particles with the actin filament in focus. This difference in contrast is consistent with an approximately 40 – 60 nm difference in height above the actin ^26^. Should myosin commonly bind and process on the side of the actin, parallel to the glass surface, no such contrast difference would have been observed.

Also, the off-axis unbound state reported in Andrecka et al.- Figure 8 has a mostly circular probability distribution. If a myosin-5a molecule started off parallel to the coverslip and rotated around the actin azimuth during its processive run, this distribution would become elliptical over the course of the run with the long axis at right angles to the actin filament axis. No such change in the distribution shape was observed. Also, as the myosin rotated around the actin the mean distance between the center of the 20 nm bead and the coverslip would vary from 10 – 25 nm above the glass surface. We would expect this to be accompanied by a change in signal contrast, however contrast of the bead shows little change between successive bound states ^26^.

For these reasons, together with those discussed in Andrecka et al. (specifically, in the section “Revealing the unbound head motion in three dimensions”), we determine that the myosin-5a in these experiments mostly binds and processively moves on top of the actin filament, perpendicular to the glass surface, with a limited range of angles bounded by the actin filament attachment to PEG-PLL/glass surface.

### Stride distribution analysis

Myosin-5a stride lengths were plotted as a histogram (Figure 2A) binned using a bin-width optimization algorithm ^34^. The binned data were then fit with a sum of seven Gaussian functions, each with the same fixed width and different means and areas, using a least-squares method. The fixed width was determined as the value (1.195 nm) that minimized the error on the total fitted area of the sum of seven Gaussian functions. The probabilities of step lengths were calculated by a least squares best fit solution to the set of simultaneous equations containing the probabilities of the underlying step lengths. The KDE plot was created using an adaptive bandwidth algorithm ^16^.

To test whether consecutive strides were independent events, all pairs of consecutive strides (629 pairs from 96 runs that totaled 725 strides) were entered into a 9×9 grid (stride lengths 20 to 36 actin subunits). The relative frequency of each stride length was used to calculate the expected number of each pair for a chi-square test of the null hypothesis that strides are independent. There was no pattern to the distribution across the grid of differences between observed and expected values. Using the 22 cells for which the expected numbers were greater than 5, p>0.05, so the null hypothesis was accepted.

### Myosin-5a cryoEM analysis

Among 1,137 myosin-5a molecules identified in 57 micrographs, we found frequent (55%) extended molecules bound to F-actin by both motor domains (Figure S3), and small proportions bound by one motor domain (9%) or bridging between two actin filaments (4%). Very few (2%) compact molecules were associated with actin, as expected from our earlier work ^35^. Remaining molecules were not attached to actin and were either extended (11%) or compact (18%).

Image processing comprised the following steps carried out within the SPIDER software package ^36^. Images were contrast inverted (protein light) and CTF corrected by phase flipping. Decorated and undecorated F-actin filaments were segmented, aligned, and classified on features within actin to generate classes of polar filaments as judged by substructural detail. Those possessing bound myosin molecules were then selected (493 molecules) into a separate data set. To measure the distance between leading and trailing head motors along the actin filament we classified the images based on features around the motor domain, thus putting leading and trailing motor domains into different classes. We then marked the position of the motor domain (by eye) in averaged images of those classes where the motor domain was clearly visible. The distance between paired motors could then be computed from the raw images. Identifying the position of the motor domain was helped because the position of the actin subunits to which they are bound was very clear in these image averages. For generating averaged images with motors with different separations, having identified F-actin polarity (above) it was then necessary to mirror invert those images of myosin-bound molecules so that when displayed with F-actin running horizontally, all the myosin molecules were above the filament. This was done by image classification based on myosin features. Then all actin-bound myosin molecules were brought into a common alignment in which the molecules were walking to the right. Histograms were created using GraphPad Prism.

For generating averaged images with motors with different separations, having identified F-actin polarity (above) it was necessary to mirror invert those images of myosin-bound molecules so that when displayed with F-actin running horizontally, all the myosin molecules were above the filament. This was done by image classification based on myosin features. Then all actin-bound myosin molecules were brought into a common alignment in which the molecules were walking to the right.

### Myosin-5a nsEM analysis

Data analysis was conducted primarily in SPIDER software ^36^. Molecules with both motors bound to actin were picked as two coordinate pairs, corresponding to both motor domain positions (n=1073 molecules). Using the coordinates, images were rotated to bring the actin horizontal with both motor domains positioned above the actin. Visual inspection of the stack of images was used to split the data into two, one subset with molecules walking right, one subset with molecules walking left. Leading heads from each dataset were subjected to reference-free alignment with restricted rotation and shift to create an aligned stack in which motor domains were superimposed. The same was done for trailing heads in each dataset. Each of the four aligned stacks was classified and divided into 20 classes per stack (= total of 80 classes) using K-means classification using a mask focused on the motor domain. A position was marked in the clearest class averages created from each dataset, corresponding to the contact point between the motor domain and actin (55/80 classes = 69% selected for defining contact points). These coordinates, along with the rotations and shifts from previous steps, were used to compute the location of the contact point in the original images and the motor-motor distance was calculated (n=566 molecules). Datasets were subdivided into classes based on motor-motor distances and averaging of these subclasses was used to validate the assignment of directionality and motor-motor distance measurement. Motor-motor distances were plotted as a histogram with adaptive bandwidth KDE (Figure 3F) using the same method as the iSCAT analysis (Figure 2A).

### Analysis of cryoEM data for the presence of CAD

CryoEM images of frozen hydrated chicken skeletal ADP-F-actin, at a resolution of 0.103 nm/pixel, were downloaded from EMPIAR dataset 11128^1^. Subunit detail was enhanced by Fourier bandpass Gaussian filtering between 50 and 20 pixels using Fiji-ImageJ 1.53. This allowed filament polarity to be assigned from the polar appearance between the crossovers (Figure S4D). Starting at the pointed (-) end of an uncluttered segment of an actin filament, consecutive crossover positions in a filament were marked by eye, using the Multipoint utility of Fiji-ImageJ, to create a dataset from n = 54 filaments. The sets of x, y coordinates generated for filament segments were further analyzed using Microsoft Excel to determine each crossover spacing and standard deviations of spacings. Distances from the starting crossover position to each subsequent crossover position were also computed. However, clutter sometimes limited the length of filament segment that could be used, with the result that crossover distances to the subsequent crossovers 1 – 4 could be measured on all 54 segments, but distances to crossovers 5, 6, 7, and 8 could be measured on only 51, 39, 28, and 18 segments respectively.

### Creating model actin filaments

Model filaments were built by sequential addition of subunits starting at the pointed (-) end, according to the following rules. The 3D coordinates of the centers of the n^th^ actin subunit in a filament aligned along the x axis are given by the following set of parametric equations:

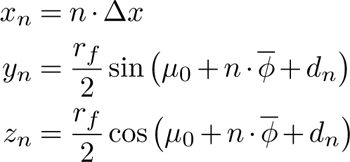

where *r_f_*is the filament radius, *φ* is the mean rotation per subunit, *d_n_* is the variable rotation per subunit, and Δ*x* is the rise per subunit. *φ* was set to −166.39°, *r_f_* to 6.5 nm, *d*_0_ to 0°, and Δ*x* to 2.75 nm. For visualization, the azimuth of the initial subunit (*µ*_0_) is set to 60°, changing this initial value equates to a rotation of the whole filament around the x axis. For each sequential subunit the value of *d_i_*was a random selection from a Gaussian distribution of values having a mean of 0° and a standard deviation of *d*. Visualization of these filaments (Figure 1B, 4A, 4B, and 5A) were created in Adobe Illustrator using the calculated coordinates as guides for actin subunit centers. Myosin motor binding site coordinates were estimated, for purposes of visualization, in a similar manner centered at *r_f_* = 6.6 nm and setting the initial azimuth value to 110°.

### Analysis of crossover distances of model filaments

A side view of each model filament was studied by only considering the x and y coordinates. The filaments were modelled as two coaxial, long-pitched sinusoids by taking alternate subunits along the helical axis. The coordinates of the intersections of these two sinusoids were calculated using the Shapely package for Python ^13^. If position error is included in the model, randomly generated noise from a Gaussian distribution with mean 0 nm and standard deviation of the position error was added to the intersection coordinates. Crossover distances were calculated by taking the Euclidean distance between adjacent crossovers.

Adjacent crossover spacings were plotted as scatter graphs (Figure S4B). To fit a regression line through this scatter, we used orthogonal distance regression (ODR) a specific case of Deming regression in which the error of variances in x and y are equal ^12, 37^. This yielded a slope of −1.00 ± 0.016. Therefore, to obtain a measure of the ellipticity of the scatter plot we computed the standard deviation of the data projected onto an axis of slope −1 (SD_-1_) and the orthogonal axis of slope +1 (SD_+1_). To do this, the (x, y) coordinates of the data points were first rotated clockwise by an angle *θ* = 45° about their mean using the following transformation matrix:

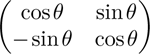

and then SD_-1_ and SD_+1_were calculated as the SD of the y and x coordinates respectively. The ratio of the two SD values was calculated as SD_-1_/SD_+1_. Variation in this ratio for given conditions was determined through 1000 iterations and plotted as a Gaussian with mean and standard deviation of the ratios in this iterative dataset (Figure S4B, inset). Additionally, variation in the ratio as a function of both CAD and picking error was determined from 25 error values, evenly spaced, from 0 – 5 nm and 50 CAD values from 0 – 10° (Figure S5). The value of any one of these three values (ratio, CAD, error) can be determined from the other two by interpolation of this dataset using a Clough-Tocher 2D interpolator function ^12^.

The SD of distance along the model filaments as a function of crossover number was determined by taking the standard deviations of the Euclidean distance between each crossover and the first crossover of a sample of filaments (Figure S4C and Figure S6A). The variance is calculated as the square of these values (Figure S6B and C). A least squares linear fit to the variance is obtained ^12^.

### Determining CAD and position error in EM data

The average SD in crossover spacing due to the presence of CAD was calculated. From inspection of a helical net projection of the actin helix (Figure S4A), it is app<u>a</u>rent from similar triangles that for a giv<u>e</u>n mean crossover spacing, *x*, as measured in the EM data, *x* / −180 = ‘6.*x* / (−180 - *φ*). Thus, the mean rotation per subunit, *φ*, can be calculated:

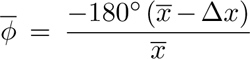

For a given RMSD variable rotation per subunit, *d*, the RMSD CAD at the n^th^ subunit, *σ_n_*, is given by:

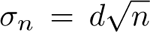

Dividing *σ_n_* by the mean crossover spacing, expressed in number of subunits, a mean change in rotation per subunit, Δ*φ* can be obtained:

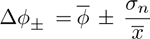

This can be converted into a change in crossover spacing by substitution of Δ*φ_±_* into the equation for *φ* above.

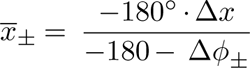

The mean of the resulting values, *x*_+_and *x_−_*, is taken as the contribution CAD makes to the crossover spacing SD value. The square of this value will give the contribution to the variance, Var_CAD_.

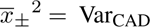

Var_CAD_ scales approximately linearly with CAD over the range 0 – 10° (Figure S6F), although deviates from linearity at higher CAD values.

The contribution of error from picking to the observed variance of crossover spacings in the EM data, Var_EM_, can thus be calculated as:

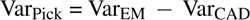

With the square root of this value giving the SD of error from picking for a crossover spacing. Since two crossover picks are made to determine a crossover spacing, the variance of each pic*_√_*k will be half that associated with the spacing measurement, and hence the SD for picking a crossover is (error from picking)/2. Through least squares fitting ^12^, an optimal CAD value can be obtained from the measured mean crossover spacing in a data set that produces a line of best fit of form (*n* Var_CAD_ + Var_Pick_) (Figure S6E-F).

### Actin disorder analysis

For an actin filament with exactly 13 subunits in 6 turns of the left-handed short-pitch helix, the rotation around that helix per subunit, φ, (−166.154° in this case) is such that 13φ= 6(−360). The difference in azimuth around the actin filament axis between the zero^th^ and the 13^th^ subunit, µ_13_ = 13φ- 6(−360). If φ has a larger value, µ_13_ will be negative. For the 11^th^ subunit, µ_11_ = 11φ– 5(−360). Generalizing for 0*^◦^ < µ_n_ <* 360*^◦^,*

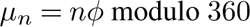

It is convenient to set the zeroth subunit to 0°, and *−*180*^◦^ < µ_n_ <* 180*^◦^*. Adjusting the equation,

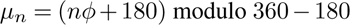

A characteristic of cumulative angular disorder is that the magnitude of disorder of a subunit is independent of the subunits around it. Therefore, the variance (σ_n_^2^) in subunit azimuth relative to a given starting subunit increases linearly with the number of subunits traversed, and thus:

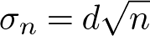

Where d is the RMSD disorder value in degrees.

Modelling an actin filament as a cylinder, a Gaussian azimuthal probability distribution of the n^th^ motor binding location will be ‘wrapped’ around the circular cross section of the cylinder. This imposes a periodic boundary condition upon the distribution. Consequently, estimated azimuthal probability distributions (Figure 5C and F) for each step length (*n*) were calculated as wrapped normal distributions, with mean *µ_n_*, using ^38^:

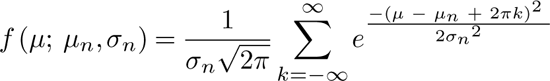

The distributions were rescaled by a common factor such that their values at an angle of µ = 0*^◦^* summed to 1 (Figure 5D and G). Optimal φ and *d* values were obtained through least-squares fitting of the values of probability of each subunit being at zero azimuth to the iSCAT values of relative frequencies of step and stride lengths.

### iSCAT dwell-time analysis

The dwell-time analysis takes account of the fact that the duration of each step along actin is determined by the rate constants of ADP release from the trail head and the subsequent ATP binding to it that triggers trail head detachment from actin. Fitting of the dwell time measurements therefore uses the formula for a two-step reaction ^39^, in which k_1_ conventionally refers to the second order ATP binding step and k_2_ to the first order ADP release step, though these events occur in the reverse order in an attached head. At the concentration of ATP used in the iSCAT assay (10 µM), the observed first order rate constants of both processes are expected to be about 10 s^-1^ as discussed previously ^26^. We therefore analyzed the dwell times using two models (Figure S6), one setting them to be equal, the other (arbitrarily) setting k_s_ to be less than k_f_. In the latter case we are not specifying which of the two enzymatic steps has the smaller rate constant. Dwell times for the A and B states were pooled into a single dataset distinguished by the stride length preceding or succeeding the A and B state respectively. Rate constants for each stride length were determined through maximum likelihood fitting by minimizing a single global negative log-likelihood function for all strides. The distribution used is a convolution of two exponentials ^40^ of the form:

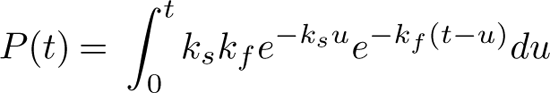

if the two rates are not equal this evaluates to:

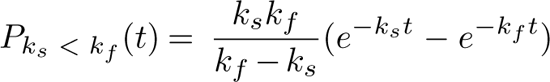

and if the two rate constants are equal, k_s_ = k_f_ = k:

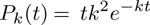

---

**Figure S1.**
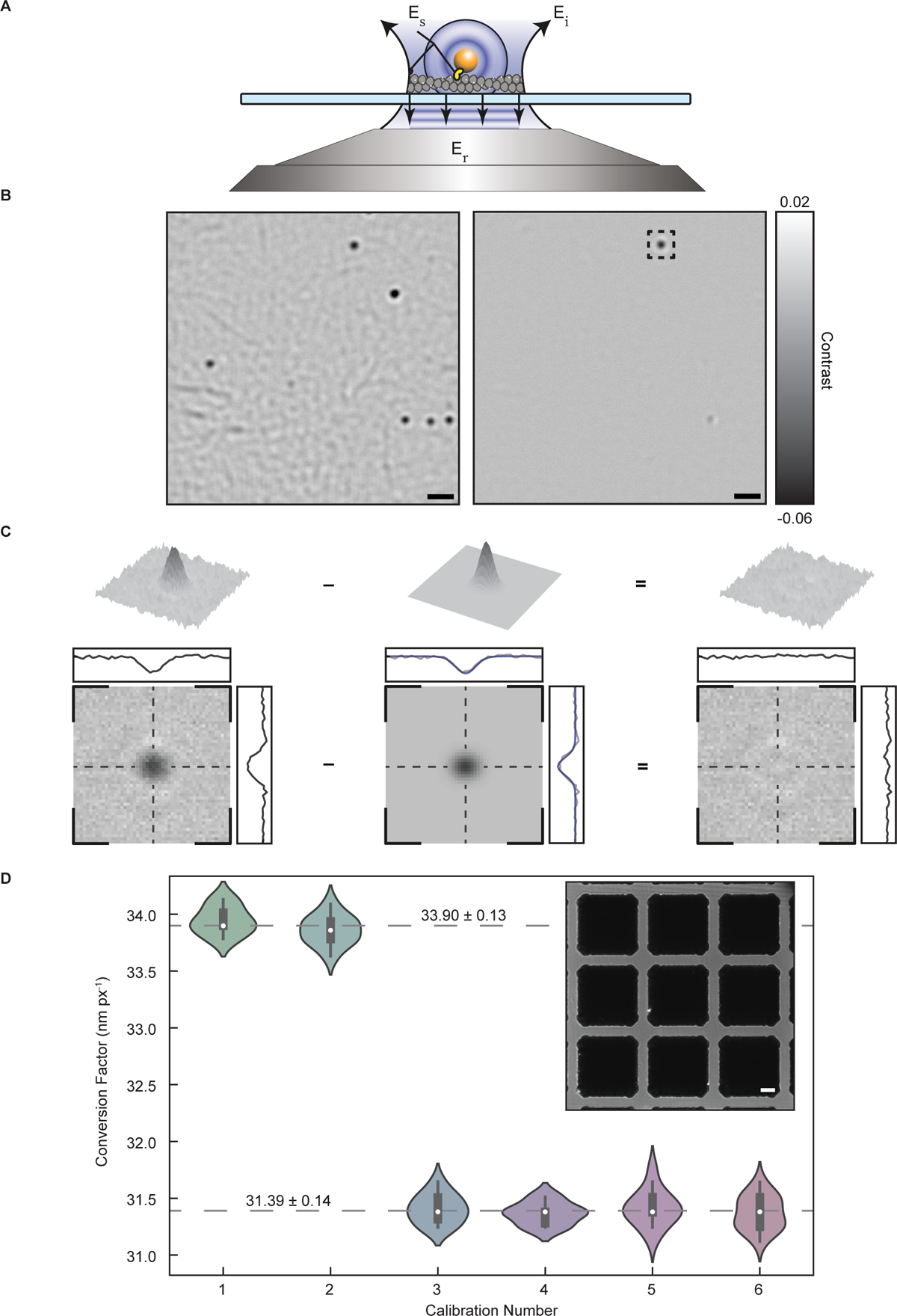
Essential principles of iSCAT microscopy. Related to. Figure 1. (A) Schematic of iSCAT showing incident, reflected and scattered light (Ei, Er, and Es respectively). A coverslip (blue) supporting actin and myosin-5 labelled with a gold bead is imaged in an inverted microscope using an oil-immersion objective (grey). (B) (Left) Raw iSCAT image of unlabeled actin network with gold-labelled myosin-5a. (Right) Image with static background subtracted, which reveals only the gold nanoparticles that have moved since the static background image was recorded. Scale bar (black) = 1 µm (C) Image of the gold nanoparticle highlighted in (B) and the 2D Gaussian fit subtracted to show the fit residual. Data are shown as both an XY plus contrast axis plot and an image. (D) Calibration of iSCAT camera pixels. Calculated conversion factor from pixels to nanometers, as determined by use of a resolution grid (Inset; scale bar (white) = 2 µm). Microscope adjustments were carried out between calibrations 2 and 3. Violin plot envelopes are kernel density estimates of the distributions with bandwidth determined by Scott’s rule of thumb. Grey dashed lines indicate two global means before and after microscope adjustment, together with standard error of mean.

**Figure S2.**
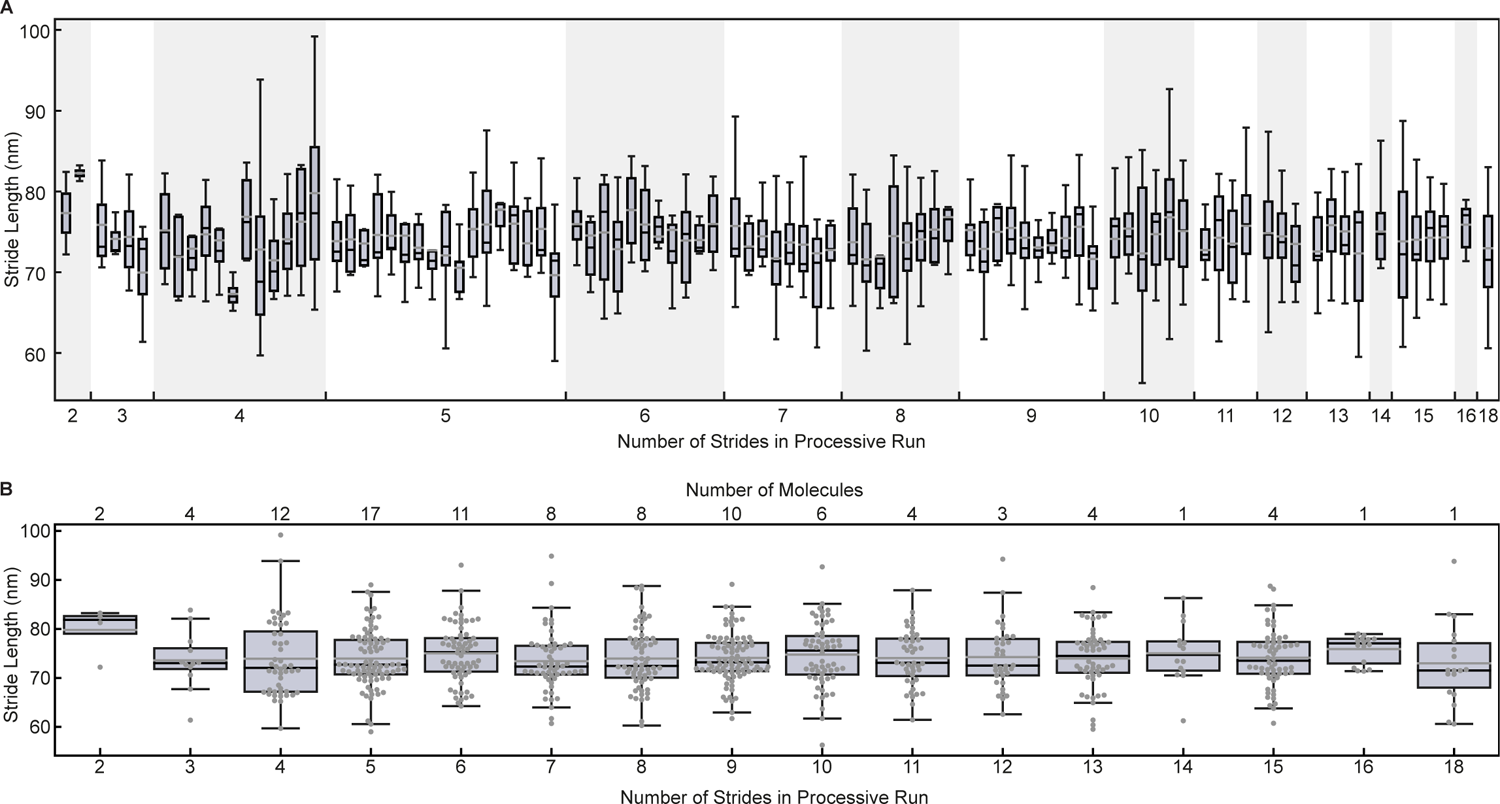
Characteristics of myosin-5a processive runs. Related to. Figure 2. (A) Distribution of stride lengths for all tracked myosin-5a molecules. Molecules have been grouped by the number of strides in the run. Box plots of lengths of strides in processive run with each box corresponding to a single myosin molecule (N = 96 molecules, 725 strides). Boxes show quartiles of the dataset with error bars extending 1.5 times the inter-quartile range. Median and mean stride lengths marked in horizontal black and grey lines respectively. (B) Stride length is independent of processive run length. Data from each group in (A) have been combined. Grey dots mark every individual stride. Boxes show quartiles of each dataset with error bars extending 1.5 times the inter quartile range. Median and mean stride lengths marked in horizontal black and grey lines respectively.

**Figure S3.**
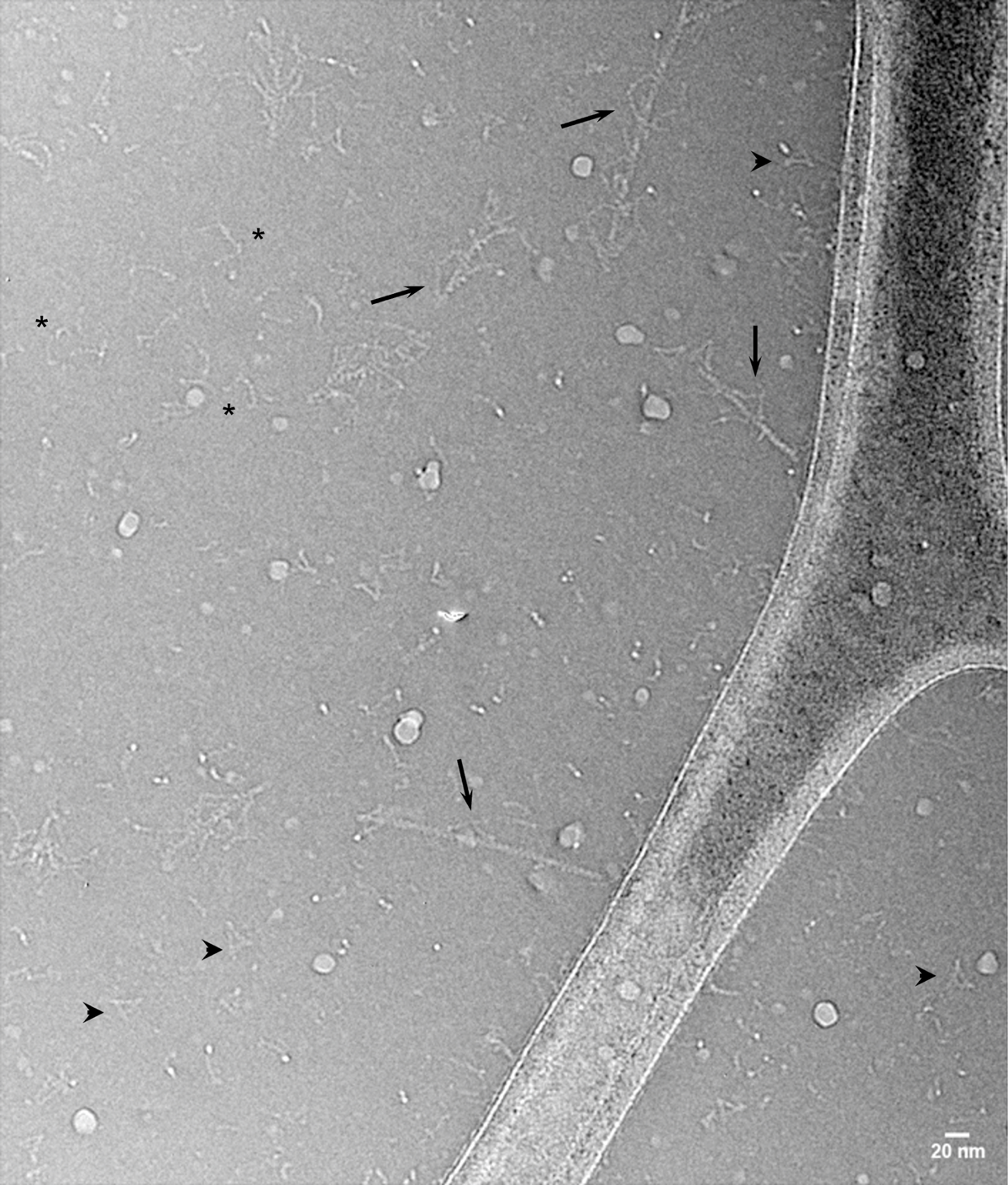
CryoEM field of full-length myosin-5a and F-actin in the presence of ATP. Related to. Figure 3. Examples of extended myosin molecules bound by both heads to F-actin (arrows). Also compact molecules (arrowheads) and extended, detached molecules (asterisks). Contrast inverted (protein white). Scale bar: 20 nm.

**Figure S4.**
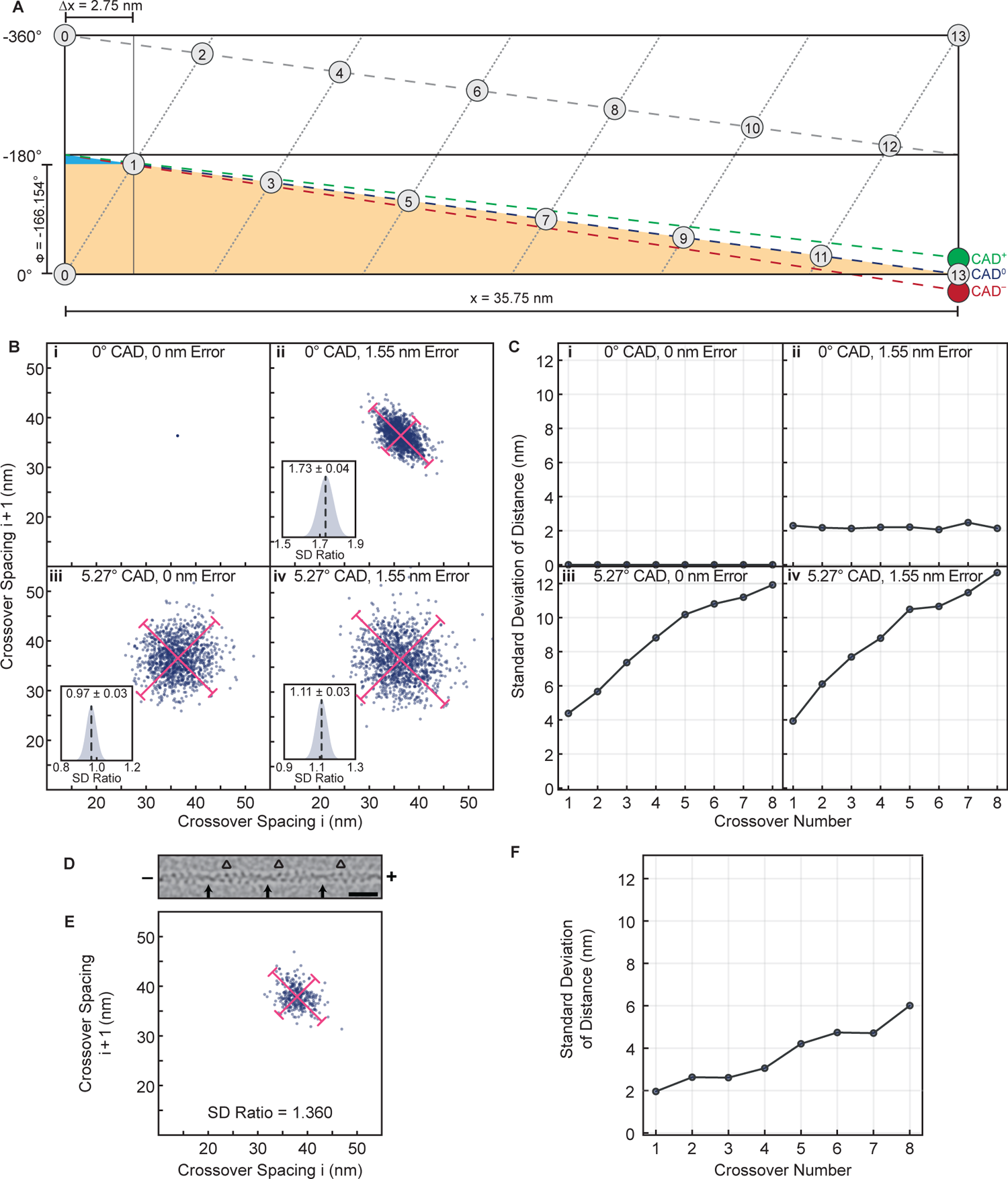
Analysis of actin filament CAD from cryoEM images. (A) Helical net representation of F-actin for deducing the relationship between rotation per actin subunit and crossover spacing. The net can be thought of as being made by dropping a hollow cylinder around an actin filament, marking the positions of the subunits on it, then cutting the cylinder lengthways and opening it out flat. The single, short-pitch, left-handed helix that passes through all the subunits is marked with short dashes; the two long-pitch, right-handed helices are marked by long dashes. Each crossover is where the long pitch helices cross the 0° and −180° lines. The net shown corresponds to the classical F-actin 13/6 helical geometry (13 subunits in 6 turns of the short-pitch helix), but the concept is general. The rotation per subunit (φ) of −166.154°, the rise per subunit (Δx) of 2.75 nm, and the crossover spacing (x) of 35.75 nm are all shown. CAD that produces a net decrease of subunit rotation over the 13 subunits (CAD^—^) is shown by the red dashed line, which can be seen to move crossover position to the left, shortening crossover spacing. CAD^+^ (green dashed line) is the complementary phenomenon. Note the two similar triangles: the larger (orange) with apexes at 0°, −180°, actin subunit 13 and the smaller (blue) with apexes at −166.154°, −180°, actin subunit 1. (B) Scatter plots of successive actin crossover spacings in simulated actin filaments (100 subunits, 1,000 filaments) with and without CAD and Gaussian error in picking crossovers. Error bars denote 1.96 * the standard deviations along lines of slope −1 and +1. Insets: Gaussian distributions of the spread in the ratio of the standard deviations along the lines of slope −1 and +1, generated from 100 repeats. (C) The standard deviation of sequential crossover distances in 1,000 filaments, under the same conditions as in (B). (D) CryoEM image of F-actin ^1^, bandpass filtered to enhance subunit detail. Arrows mark position of crossovers. Triangles mark the position of triangles formed by three bold actin subunits on the barbed (+) side of each crossover that allows assignment of filament polarity. Scale bar: 20 nm. (E) Scatter plot of crossover spacing (i) against the neighboring crossover spacing towards the barbed end (i+1). (F) Measurement of the spread (SD) of values among a set of actin filaments, of the distance along the actin of a series of 8 consecutive crossovers away from a starting crossover.

**Figure S5.**
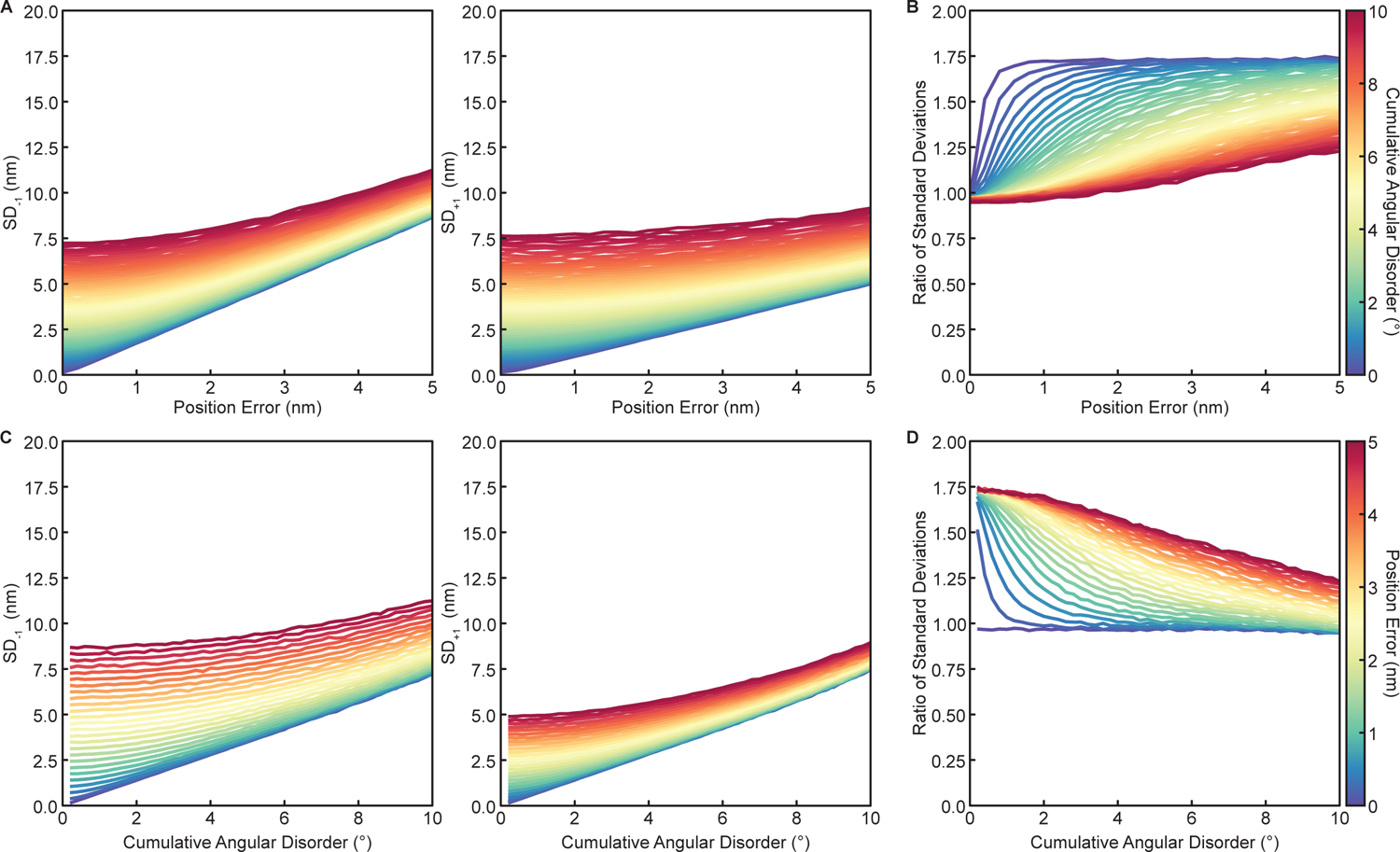
Actin filament CAD simulation of consecutive paired crossover distance. Related to Figure S4. (A) Standard deviation of simulated consecutive pair crossover scatter plots (1000 subunits, 1000 filaments) along lines of slope −1 and +1 (SD-1, SD+1) as a function of error from picking crossovers in determining the actin crossover spacings. The hue scales with magnitude of CAD. (B) The ratio SD-1 / SD+1 as a function of picking error. (C) and (D) The same simulated dataset transposed to show CAD along the x-axis and error from picking scaling as hue.

**Figure S6.**
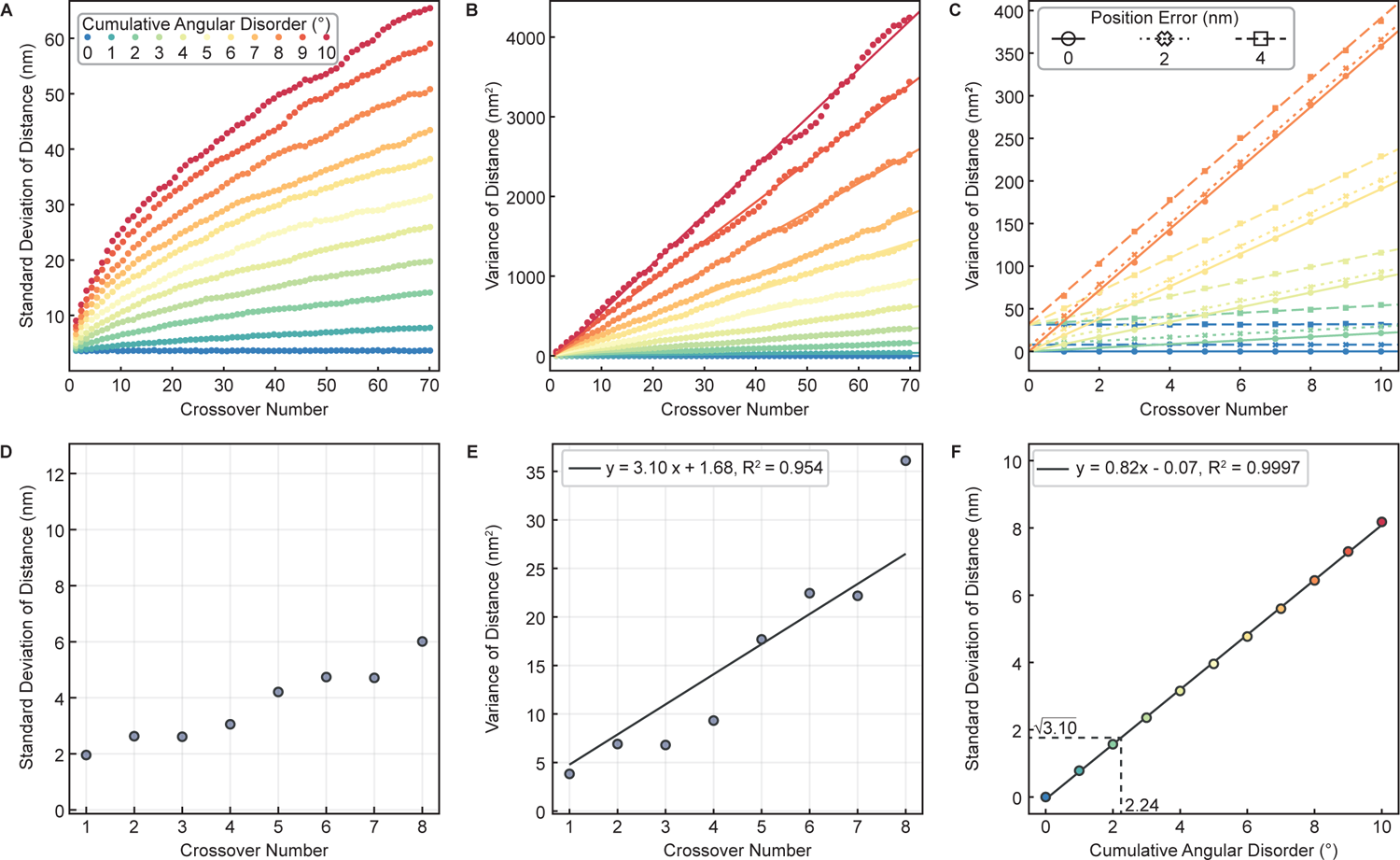
Actin filament CAD simulation of sequential crossover distance. Related to Figure S4. (A) Simulated standard deviations and (B) variances of sequential crossover distances extended to 70 crossovers per filament. The hue scales with cumulative angular disorder (CAD). Linear fits to the variances all have r^2^ 0.992. (C) Variation of gradient with picking error (10000 filaments). There is minimal change to the gradient with position errors of 0 nm (circles), 2 nm (crosses), and 4 nm (squares). Note that the offset caused by position error is given by 2(position error)^2^ (so is 8 nm^2^ and 32 nm^2^, respectively), and can be measured as the intercept on the variance axis. (D) Standard deviation plot and (E) variance plot for the cryo-EM data from F-actin ^1^ Variance plot has line resulting from least squares fitting to estimate both CAD and measurement error (F) Calculated standard deviation of the first crossover as a function of CAD. A linear fit (R^2^ = 0.9997) is shown. Interpolation of the cryoEM data at a gradient value of *^√^*3.10 as determined in (E) returns a CAD of 2.24 *±* 0.05° for the cryo-EM data.

**Figure S7.**
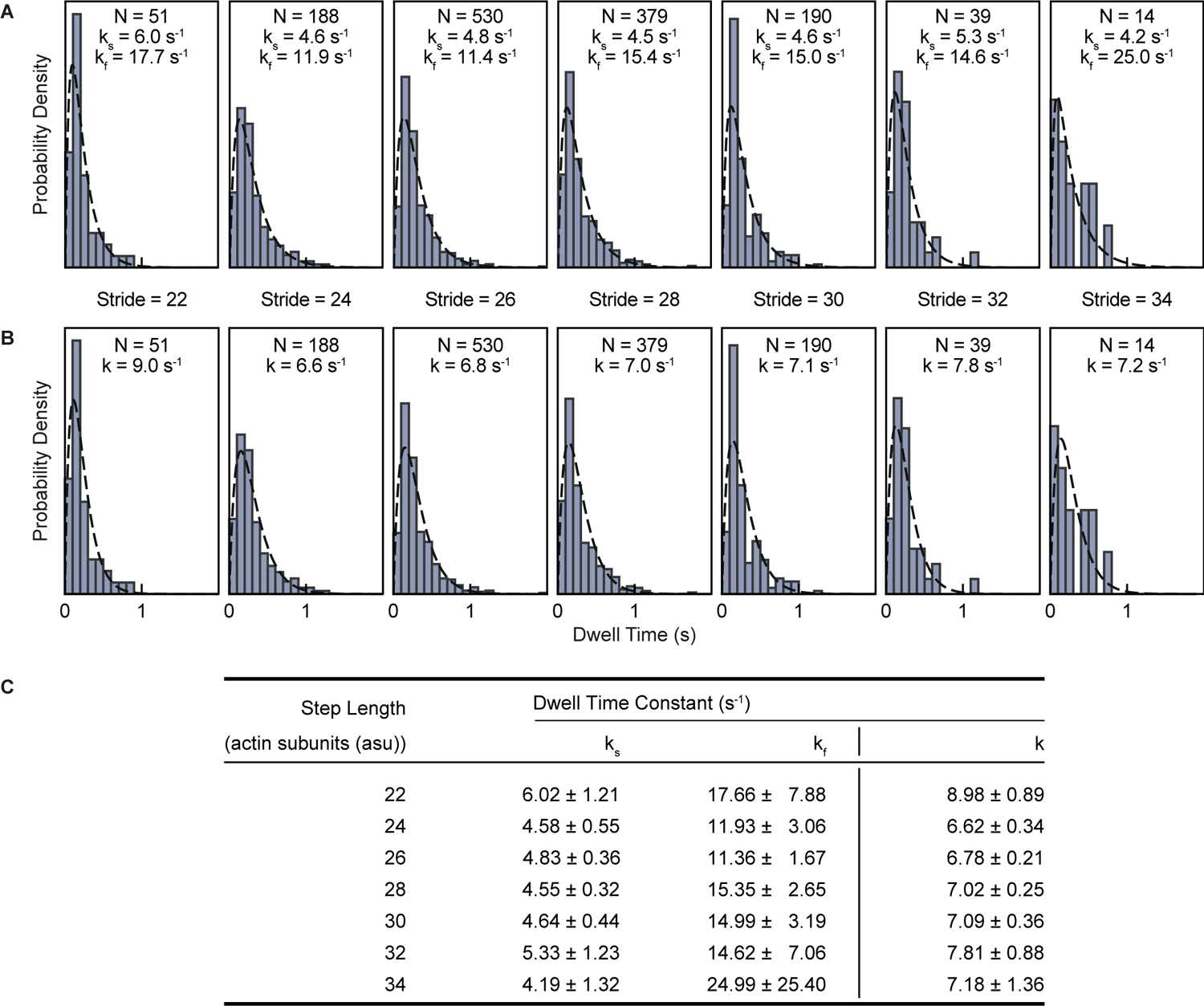
Fitting of trailing head dwell times to a two-step reaction sequence. Related to. Figure 6. Panels show dwell time distributions (bars) and results of a global maximum likelihood fitting (dashed lines) to a convolution of two exponentials. (A) Rate constants assumed to be different. k_s_ is set to be lower than k_f_ and can therefore correspond to either the rate constant for ADP release or of the subsequent ATP binding step. (B) Rate constants assumed to be the same. (C) Dwell time rate constants with error estimates are tabulated from a global maximum likelihood fitting of the dwell time data to a convolution of two exponentials when the two rate constants are allowed to differ and when they are constrained to be the same.

## Notes

### Competing Interest Statement

The authors have declared no competing interest.

